# Translational reading frame determines the pathogenicity of C-terminal frameshift deletions in MeCP2: an alternative therapeutic approach

**DOI:** 10.1101/2025.08.31.673354

**Authors:** Jacky Guy, Elena Hein, Beatrice Alexander-Howden, Timur von Bock und Polach, Tricia Mathieson, Benjamin P. Kleinstiver, Huda Zoghbi, Adrian Bird

## Abstract

Mutations in the *MECP2* gene cause the severe neurological disorder Rett syndrome. A cluster of frameshift-causing C-terminal deletions (CTDs) lead to loss of ∼100 amino acids at the C-terminus of the MeCP2 protein, and account for approximately 10% of RTT-causing mutations. The pathogenicity of C-terminal deletions (CTDs) is unexpected, as this C-terminal domain is non-essential in mice. Utilising databases of pathogenic and benign human *MECP2* mutations, we find that some individuals with apparently typical CTDs do not exhibit Rett syndrome, confirming that C-terminal truncations are not intrinsically pathogenic. Using human DNA sequence data and mouse models, we demonstrate that pathogenicity results from a drastic reduction in MeCP2 levels and is determined by the presence of the short amino acid motif proline-proline-stop (-PPX) at the C-terminus, which results from a shift to the +2 reading frame. Individuals with CTDs that shift to the +1 frame avoid this motif and do not develop Rett syndrome. Mutating the stop codon of the PPX motif to tryptophan rescues MeCP2 expression and RTT-like phenotypes in a CTD mouse model. Finally, we demonstrate that an adenine base editor can efficiently introduce this tryptophan substitution in cultured cells. Overall, our findings uncover a simple and reliable prognostic distinction between benign and pathogenic CTDs and provide proof-of-concept for an editing strategy that potentially corrects all disease-causing CTD mutations.

## Introduction

Rett syndrome (RTT, OMIM #312750) is a severe neurological disorder caused by heterozygous mutations in the X-linked *MECP2* gene (Amir et al., 1999). RTT mainly affects girls, with an occurrence of around 1:10,000 births. At approximately 18 months of age normal development stagnates and acquired skills such as speech and purposeful hand use are lost. RTT is a devastating condition, affecting speech, feeding, digestion, breathing, heart rhythm, ambulation, hand use and susceptibility to seizures (Hagberg et al., 1983; Hagberg, 2002). MeCP2 protein is found in the nucleus of all cell types but is most highly expressed in neurons (Lewis et al., 1992; Ross et al., 2016; Skene et al., 2010) where it has been implicated in a range of functions, including splicing, microRNA processing, transcriptional activation and repression (Lyst and Bird, 2015). Analysis of patient mutations and experimental data from mouse models support a role in transcriptional repression (Gabel et al., 2015; Kinde et al., 2016) although its loss leads to both increases and decreases in gene expression (Bajikar et al., 2025). Two functional domains have been characterised in MECP2: the methyl-binding domain (MBD), through which MeCP2 binds to 5-methyl cytosine in DNA (Nan et al., 1993), and an NCoR interacting domain (NID) (Lyst et al., 2013; Nan et al., 1998). Through the NID, MeCP2 binds to TBL1X/TBL1XR1, subunits of the nuclear corepressor complex NCoR (Kruusvee et al., 2017; Lyst et al., 2013), indicating that MeCP2 acts as a bridge, bringing the corepressor complex to methylated sites in the genome.

RTT can be caused by a range of inactivating mutations, including large deletions, missense and nonsense point mutations, and frameshifting deletions and insertions resulting in a truncated protein. Missense RTT mutations cluster within the MBD and NID and, with a few exceptions (Guy et al., 2018), are not found elsewhere in the protein. On the contrary, neutral polymorphisms found in the gnomAD database tend to avoid the two functional domains but are commonly found throughout the rest of the protein. Alpha missense pathogenicity scores correlate with the presence of RTT mutations and absence of gnomAD variants in the *MECP2* coding sequence (Figure 1A). Using knock-in mouse models it has been shown that the parts of MeCP2 outside the MBD and NID are non-essential, resulting in a “minigene”, ΔNIC, that retains MeCP2 function (Tillotson et al., 2017) (Figure 1A).

**Figure 1.**
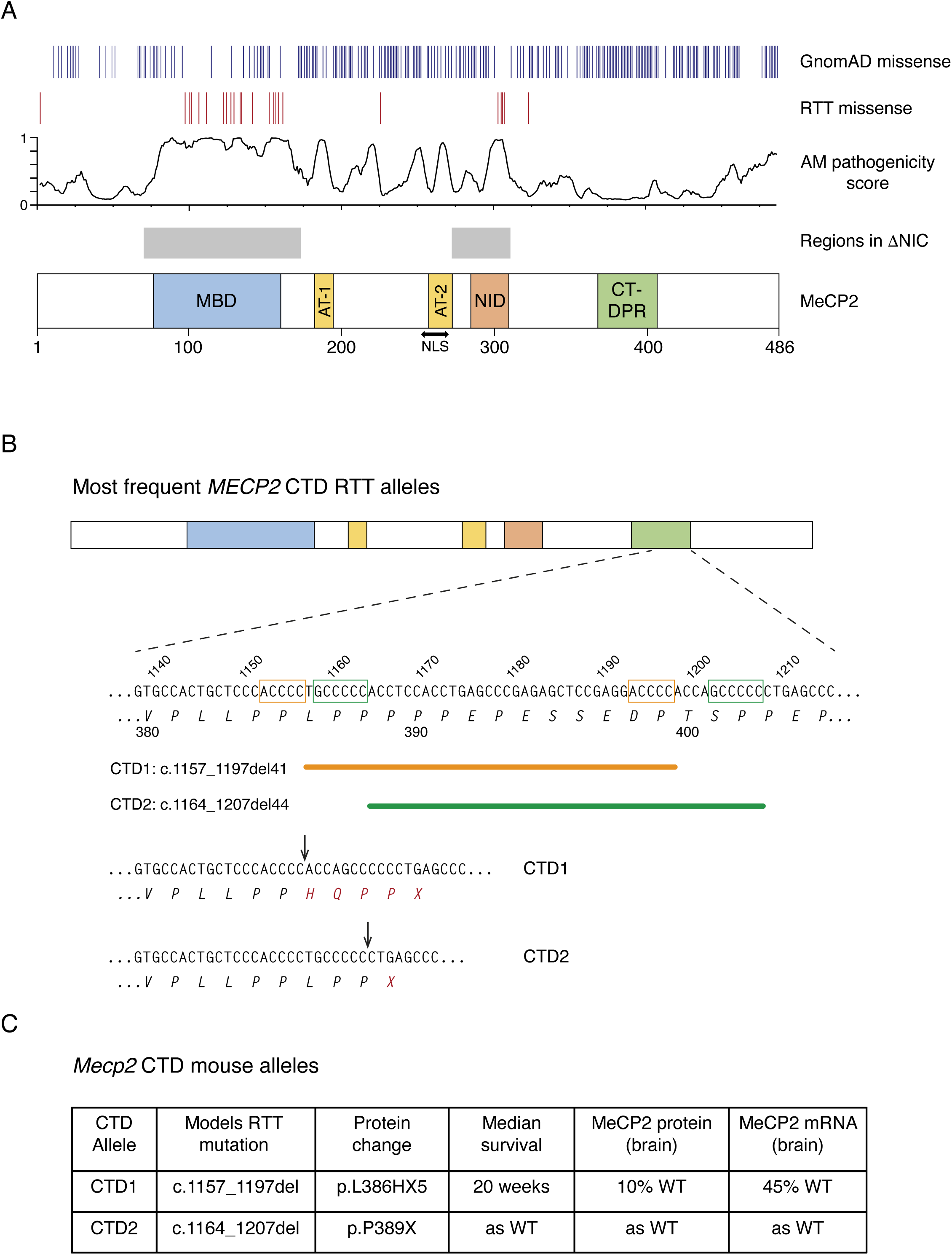
Frameshifting deletions in the C-terminal domain of MeCP2 cause Rett syndrome in humans and RTT-like phenotypes in mice. (**A**) A schematic representation of human MeCP2 protein showing hemizygous missense mutations found in gnomAD v4.1.0; *de novo* Classical RTT mutations found in RettBASE; Alpha Missense pathogenicity score; regions found in an MeCP2 “minigene” (ΔNIC) and domains described in the literature: methyl-binding domain (MBD), AT-hooks (AT-1 and AT-2), nuclear localization signal (NLS), NCoR interacting domain (NID) and the C-terminal deletion-prone region (CT-DPR). (**B**) DNA and amino acid sequence found in the CT-DPR, numbered according to transcript ENST00000303391.11, e2 isoform. The two most common RTT deletions are shown, referred to as CTD1 and 2 for brevity. Microhomologies believed to recombine to cause the deletions are marked on the DNA sequence as orange and green boxes respectively, and the deleted sequences as corresponding lines below. The C-terminal amino acid sequences of CTD1 and CTD2 are shown, with the points of frameshift marked with arrows. (**C**) Summary of CTD1 and CTD2 knock-in mouse models.

This study concerns C-terminal deletion (CTD) mutations, which comprise about 10% of all RTT-causing mutations, comparable to the most common missense and nonsense mutations (Bebbington et al., 2010; Cuddapah et al., 2014; Smeets et al., 2005). CTDs occur in a ‘deletion-prone region’ (CT-DPR) between approximately c.1110 and c.1210 (numbering according to transcript ENST00000303391.11, NM_004992, e2 isoform), and lead to truncated MeCP2 protein lacking around 100 amino acids from the C-terminus, with a missense tail of variable length (Bebbington et al., 2010). More than 50 different CTDs can be found in the database of RTT mutations, RettBASE (Christodoulou et al., 2003; Krishnaraj et al., 2017), ranging in frequency from single occurrences to over 40 individuals with the most common deletions (Figure 1B). Taken as a class, CTDs are at the milder end of the RTT clinical severity spectrum, comparable to the common R306C and R294X point mutations (Bebbington et al., 2010; Cuddapah et al., 2014). Our previous CTD knock-in mouse models (Figure 1C) revealed that the C-terminal region removed by the frameshift is not required for MeCP2 function, as mice that expressed normal levels of the truncated protein (CTD2) were not detectably different from wildtype. Instead, the disease phenotype results from a drastic reduction in the amount of MeCP2 protein present in the brain, as seen in CTD1 mice (Guy et al., 2018). Intriguingly, the gnomAD database (Chen et al., 2024; Gudmundsson et al., 2022), which lists DNA sequence polymorphisms that are unlikely to be severely pathogenic in humans, includes several individuals exhibiting what appear to be typical frameshifting CTDs. Here we uncover the genetic basis of pathogenic and non-pathogenic CTDs by demonstrating that the difference is determined by the amino acid sequence at the extreme C-terminus of the truncated protein, specifically the presence of a –PPX motif. Our findings have important prognostic implications in predicting the clinical outcomes of *MECP2* mutations. Using the knowledge gained from studying both pathogenic and benign CTD mutations we have designed and tested a strategy to edit RTT CTD mutations in the genome using adenine base editing (ABE) technology, changing the C-terminal amino acid sequence and restoring WT levels of MeCP2 protein. This provides a potential therapeutic route which could treat the entirety of this diverse class of mutations using a single ABE/guide RNA combination.

## Results

### The MeCP2 deletion-prone region shows a high level of neutral variation in the human population

To investigate the effect of C-terminal amino acid sequence on the level of CTD MeCP2 we utilised the wealth of available human mutation data. RTT mutations were collated in RettBASE (Christodoulou et al., 2003; Krishnaraj et al., 2017). Although RettBASE is no longer publicly available, the data it contained was downloaded and used in this study (Supplementary Tables 1-4). RettBASE data has now been incorporated into ClinVar (Landrum et al., 2016) in an abridged form. In addition, the ExAC and gnomAD databases hold a large set of genetic variants found within the general population (Chen et al., 2024; Lek et al., 2016). Individuals who contributed to the database are not all without clinical problems, but the inclusion of a variant suggests that it is tolerated and can be classed as neutral. It should be noted that as *MECP2* is an X-linked gene, it is possible that pathogenic variants could be present in healthy females due to extreme skewing of X-inactivation towards that allele. Indeed, family pedigrees with RTT mutations in multiple generations are possible due to this phenomenon (Augenstein et al., 2009; Ravn et al., 2011) and were used to link mutations in *MECP2* to RTT (Amir et al., 1999). On the other hand, the presence of a mutation in a hemizygous male in gnomAD strongly indicates that it is a benign variant that will not cause RTT.

GnomAD data shows that missense variants are extremely common in the C-terminal deletion-prone region (CT-DPR). Between p.370 and p.405 (c.1108 and c.1215) almost every amino acid is tolerant of change, many with multiple alternative substitutions (Fig. 2 SF1A, Supplementary Table 5). Several of these variants are present in hundreds or thousands of individuals, with E397K represented over 5000 times. E397K is so common in the general population that it occurs in the homozygous state in 11 females, and was initially classed as a RTT-causing mutation (Yusufzai and Wolffe, 2000) due to its recurrent appearance in RTT patients who were later found to have additional pathogenic *MECP2* mutations (Moncla et al., 2002; Wan et al., 1999). Also of note, many apparently healthy individuals display in-frame deletions that remove segments of the C-terminal domain (Figure 2A, Fig. 2 SF1B, Supplementary Table 6). These findings strongly suggest that the amino acid sequence of this region is not important for human MeCP2 function, in agreement with phenotypic data from a number of mouse models (Guy et al., 2018; Tillotson et al., 2017). The frequency of benign missense mutations in the CT-DPR contrasts strongly with the MBD, where the protein sequence is known to be critical for MeCP2 function (Fig. 2 SF1C, Supplementary Table 7). This difference is also apparent in the AlphaMissense pathogenicity scores across these regions of the human protein (Figure 1A).

**Figure 2.**
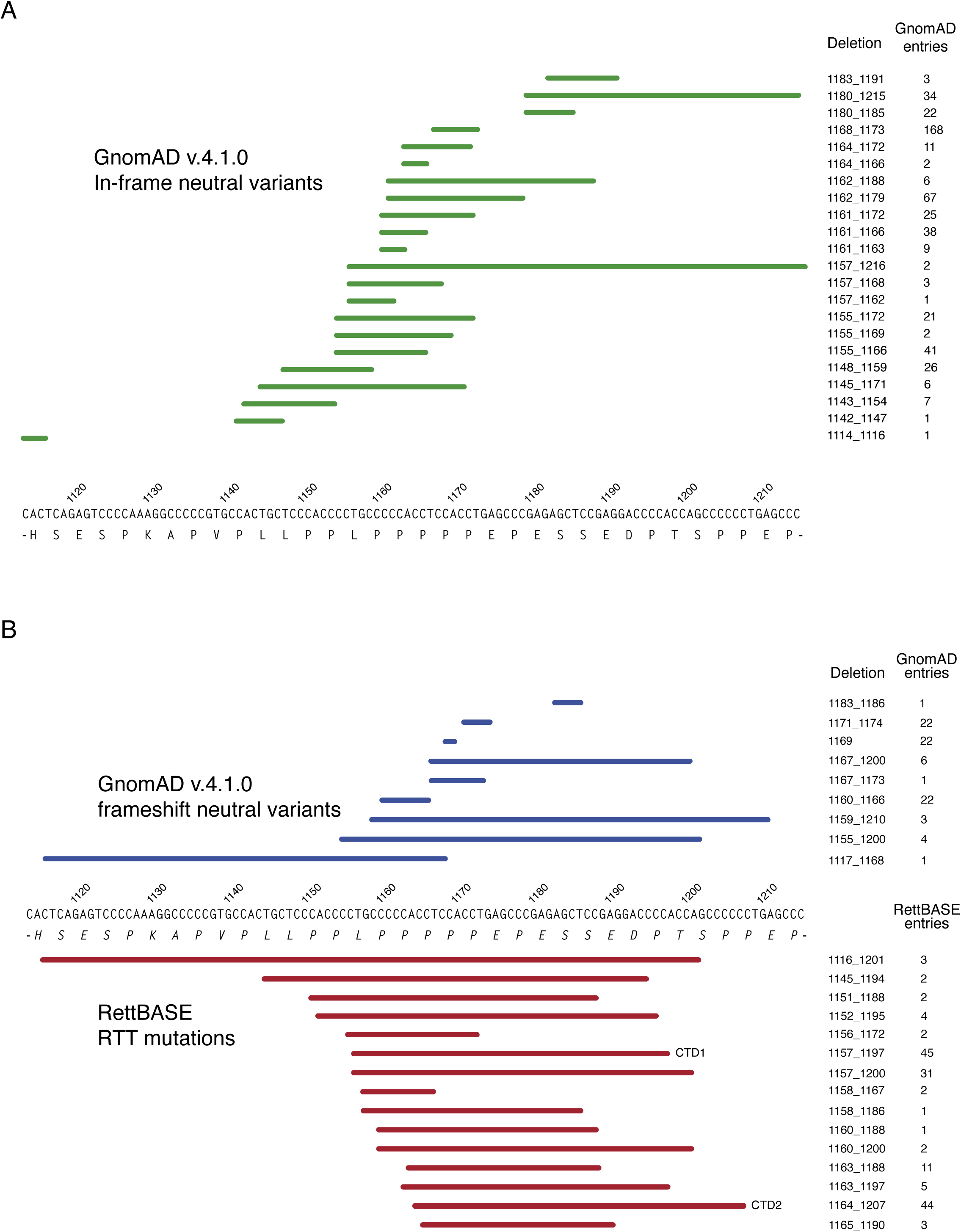
Genomic deletions in the CT-DPR of *MECP2* (high confidence sets). (**A**) All hemizygous in-frame deletions in the region found in gnomAD v4.1.0 are indicated with green lines above the DNA sequence. The deletion co-ordinates (numbered according to ENST00000303391.11) and number of individuals with each deletion are shown alongside. (**B**) Frameshifting deletions in the CT-DPR. Above the genomic sequence, hemizygous deletions from gnomAD v4.1.0 are shown in blue. Deletions from RettBASE are shown below in red, with each deletion found in at least one individual with a *de novo* mutation and a diagnosis of classical RTT. Coordinates and number of individuals with each mutation are shown, with the two most common RTT mutations, CTD1 and CTD2 indicated.

### MeCP2 C-terminal frameshifting deletions may be pathogenic or benign

Intriguingly, a substantial number of individuals in gnomAD, including hemizygous males, have frameshifting deletions that result in truncated MeCP2 protein similar to that found in RTT patients (Supplementary Table 8). To ensure that pathogenic RTT and non-pathogenic gnomAD mutations were reliably annotated, only “high confidence” sets of mutations that conformed to stringent criteria were selected for further examination. We stipulated that selected RTT mutations should be associated with a diagnosis of classical Rett syndrome and be absent in both parents (*de* novo) in at least one individual with that mutation (Supplementary Table 4). GnomAD variants were considered when present in at least one hemizygous male, to exclude the possibility of a pathogenic mutation being masked by skewing of X-inactivation in some heterozygous females (Supplementary Table 8). A comparison of these high confidence sets of benign frameshifts and pathogenic RTT CTD mutations does not reveal any obvious difference in the location of the frameshifts, with similar truncated proteins resulting in very different clinical outcomes (Figure 2B). Our previous study using CTD knock-in mice implied that the detailed amino acid sequence close to the C-terminus of CTD mutants can affect the level of MeCP2 (Fig. 2 SF2). We therefore compared the C-termini of our sets of RTT and gnomAD CTD alleles from the deletion site to the stop codon.

We found that all high confidence benign frameshifts ended in the codons –SPRTX due to a shift from the wildtype 0 frame to the +1 frame, caused by a deletion of 3n+1 nucleotides. Almost all +1 frameshifts in this region generate this ending, regardless of the exact position of the deletion. On the other hand, all but one of the high confidence pathogenic CTDs involved a shift to the +2 frame (3n+2 nucleotides deleted) resulting in most cases in the ending –HQPPX (Figure 3A, B). For both frameshifts the sequence prior to the extreme C-terminus varies depending on the size and location of the deletion. By comparing pathogenic with benign frameshifting deletions, we uncovered a striking difference between their C-terminal amino acid sequences. RTT-causing CTDs shifted to the +2 frame to give a common ending –PPX (with one exception, c.1158_1167del, CTD3; see below), whereas all benign CTDs shifted to the +1 frame to end in –SPRT (Figure 3B).

**Figure 3.**
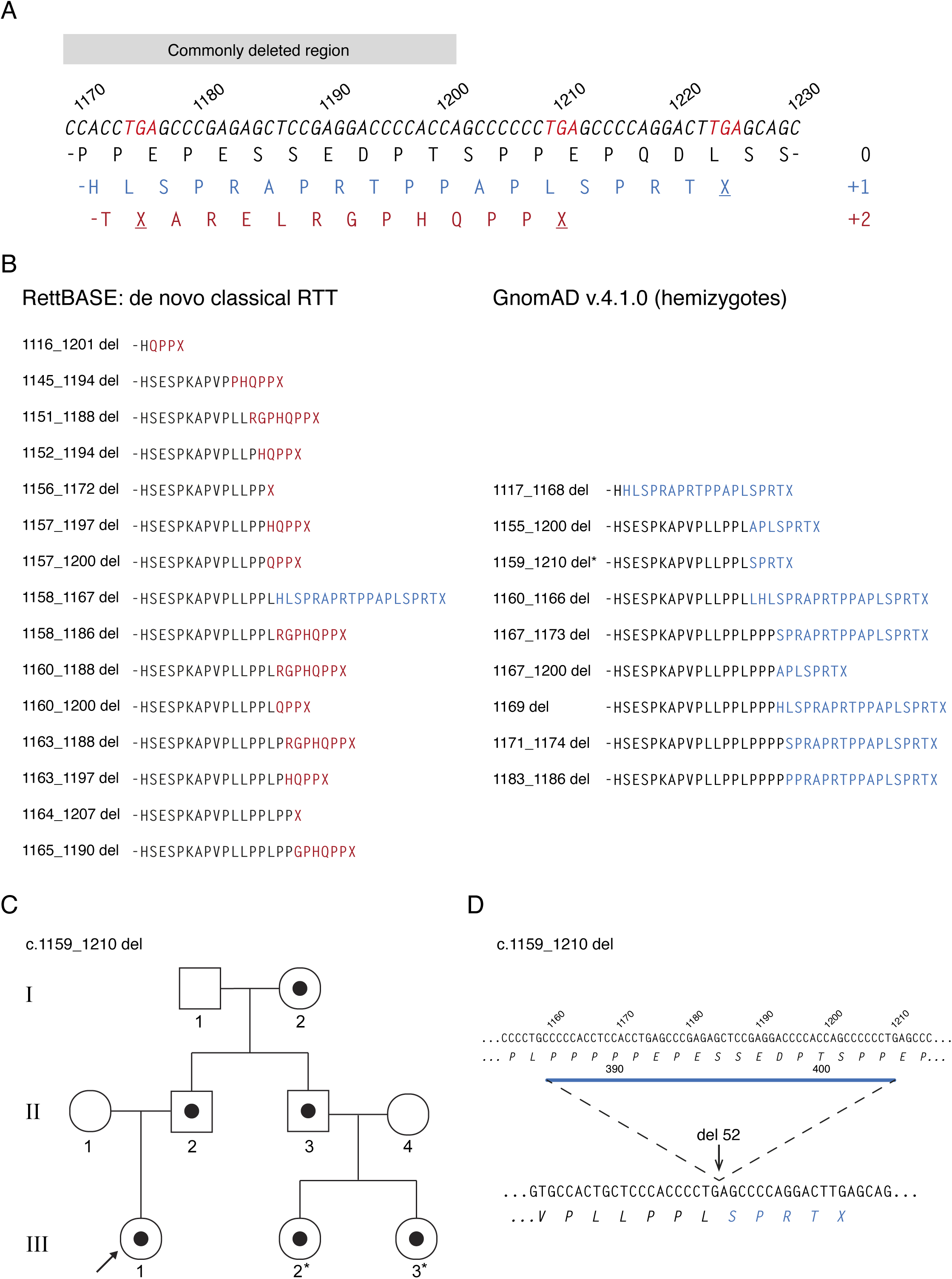
C-terminal amino acid sequences of pathogenic and benign MeCP2 CTDs (high confidence sets). (**A**) The three possible reading frames after frame shifts in the CT-DPR. WT genomic sequence is shown, with all possible stop codons in red. The amino acid sequence of the WT reading frame (0) is shown in black, with +1 frame in blue and +2 in red. (**B**) C-terminal amino acid sequences of frameshifting deletions shown in Figure 2. Sequence after frame shift is shown in blue (+1 frame) or red (+2 frame). (**C**) Family pedigree showing three generations from a family with a c.1159_1210 del *MECP2* mutation (black filled symbols). (**D**) Genomic DNA sequence and amino acid sequence showing c.1159_1210 deletion site and molecular consequences.

We expanded our enquiry to look at all CTDs in gnomAD, including heterozygotes. The most recent gnomAD release, v.4.1.0, contains sequence data from over 800,000 individuals, with a total of 103 *MECP2* CTD alleles carrying 27 different mutations (Supplementary Table 8). In line with the expectation that the pathogenic frameshift would be severely under-represented in the general population, we observed 96 individuals with 21 different alleles that were shifted to the “benign” +1 frame, whereas only 7 individuals had the potentially pathogenic +2 frame (Fig. 3 SF1A). On closer examination, all but two of the +2 frameshifts avoided the –PPX ending due to deletions which ended prior to an earlier stop codon (Fig. 3 SF1B). The two exceptional mutations were each found in a single heterozygous female individual, with one carrying the recurring c.1164_1207del RTT allele (CTD2). A possible explanation for the rare presence of these pathogenic mutations, so far untested, is that skewed X-inactivation masks their deleterious effect.

In contrast to the predominance of +1 frameshifts found in gnomAD, when all CT-DPR frameshifts in RettBase were considered, +2 frameshifts were present in a large majority of RettBASE entries (Fig. 3 SF2A, Supplementary Table 2). Once individuals without a RTT diagnosis had been excluded, there were 25 cases with +1 and 185 with +2 frameshifts (Fig.3 SF2B).

### A case study illustrating the importance of differentiating benign versus pathogenic C-terminal deletions

A pregnant woman approached one of us because prenatal testing revealed an *MECP2* c.1159_1210 deletion in the female foetus. The diagnostic company indicated this is a pathogenic deletion. The deletion was present in the unaffected father which inspired the family to reach for a second opinion. The deletion was in 100% of the father’s cells and a detailed interview with a focus on neuropsychiatric history revealed a healthy adult male. To ensure that his healthy phenotype is not due to a second site suppressor mutation, the DNA of his healthy mother and brother was analysed. Both carried the same CTD. Of note, the brother who carries the deletion has two healthy daughters who are expected to be obligatory carriers of the mutation (Figure 3C). Lastly, the infant was born and continues to be healthy at ten months of age. According to gnomAD, identical mutations have been found in 3 further individuals, including one hemizygous male (Figure 2B, Supplementary Table 8). In agreement with our prior analysis, this mutation results in a switch to the +1 reading frame leading to a benign deletion with an –SPRTX ending (Figure 3D).

### Testing the effect of the codons adjacent to the translational stop on MeCP2 abundance and function

Our analysis of human *MECP2* DNA sequences with radically different clinical symptoms suggested that pathogenicity is determined by codons immediately adjacent to the stop codon. Specifically, the ending –SPRTX is associated with almost all benign human CTDs that involve frameshifts. Unexpectedly this ending is also reported in one of the stringent set of RTT CTDs, c.1158_1167del (CTD3)(Bienvenu et al., 2000; Philippe et al., 2006). Our hypothesis predicted that this mutation had mistakenly been classified as pathogenic and is in fact benign. To test this prediction, we created a CTD3 knock-in mouse allele (Fig. 3 SF3A). The resulting male hemizygous CTD3 mice appeared like WT littermates, did not develop RTT-like phenotypes (Fig. 3 SF3B, C) and had 100% survival at one year. CTD3 MeCP2 protein and RNA were expressed at WT levels in hemizygous mouse brain (Fig. 3 SF3D-F). In contrast, CTD1 mice modelling the c.1157_1197 RTT mutation developed RTT-like phenotypes, had a median survival of 20 weeks and had reduced levels of MeCP2 protein and mRNA in brain, as previously described (Guy et al., 2018). Thus, despite the CTD3 frameshift occurring only one nucleotide downstream of that in the RTT model CTD1, CTD3 hemizygous mice did not display phenotypic defects, strongly suggesting that the CTD3 mutation is benign. The result confirms our expectation that the +1 frame ending –SPRTX is not pathogenic.

These findings conflict with the presence of two RettBASE entries found in the high confidence CTD set (Figure 2B, 3B). Both cases presented a *de novo* c.1158_1167del mutation (Bienvenu et al., 2000; Philippe et al., 2006). Our findings predict that these two exceptions are in fact benign mutations and therefore not responsible for the observed clinical phenotype. It would be interesting to revisit these cases to identify alternative causal mutations, either within the *MECP2* gene or elsewhere.

Our previous modelling of two recurrent RTT-causing CTD mutations, c.1157_1197del (CTD1) and c.1164_1207del (CTD2) (Guy et al., 2018), already pointed to C-terminal codons within the truncated transcript as the root cause of MeCP2 deficiency. Building on this information, we found that although both protein and mRNA were consistently reduced in CTD1 mouse brain, the primary transcript was produced at WT levels (Fig.3 SF3F), pointing to a post-transcriptional mechanism for MeCP2 loss.

There is considerable evidence that consecutive prolines within a messenger RNA are associated with increased stalling of the ribosome during translational elongation, both in pro– and eukaryotes. Indeed, known translation factors function to circumvent this problem (Gutierrez et al., 2013; Huter et al., 2017; Schuller et al., 2017). Although less well studied, there is also evidence that the presence of two or more prolines preceding a stop codon may also interfere with translational termination (Hayes et al., 2002; Janzen et al., 2002; Matheisl et al., 2015). Our findings suggest that this may also be the case with RTT-causing CTDs. The presence of two or more prolines immediately preceding the stop codon interferes with translational termination, thereby triggering a process similar to nonsense mediated decay (NMD). This process would differ from canonical NMD in that the premature stop codon resulting from the frameshift occurs in the final exon of *MECP2,* and is also dependent on the amino acid sequence immediately before the stop codon, rather than its location relative to splice junctions. Salvage of the stalled ribosome would result in the loss of both MeCP2 protein and messenger RNA, with the amounts of primary transcript unaffected, as is the case in CTD1 mouse brain (Fig. 3, SF3F).

### A genome editing strategy for the CTD class of RTT mutations

Despite their highly variable start and end points, our findings point to common amino acid sequence features shared by all CTDs that give rise to RTT. We used this knowledge to devise an editing strategy that would apply broadly across this mutation category. Current trials of genetic therapies to treat RTT involve delivery of *MECP2* gene constructs (ClinicalTrials.gov IDs NCT05606614, NCT05898620) using Adeno-Associated Viral (AAV) vectors. A disadvantage of this approach is that the dosage to neurons is variable and must not stray into over-expression, which is toxic (Collins et al., 2004; Gadalla et al., 2017; Koerner et al., 2018; Ramocki et al., 2010; Sinnett et al., 2017). Whilst regulatory strategies to prevent MeCP2 overexpression have been devised (Ross et al., 2025; Sadhu et al., 2023), an attractive alternative approach is to edit the endogenous *MECP2* gene so that the mutational defect is permanently corrected. This should result in WT expression levels due to the preservation of all endogenous regulatory elements. Several of the common RTT missense mutations can potentially be reverted to WT using adenine or cytosine base editors (ABEs or CBEs) (Gaudelli et al., 2017; Komor et al., 2016) but correcting the more complex CTDs would require Prime editing (Anzalone et al., 2019; Chen and Liu, 2023), which is currently less efficient than base editing and would require thorough optimisation of a specific prime editing guide RNA (pegRNA) for the correction of each mutation. Knowing that the C-terminus is not necessary for MeCP2 function and that the –PPX ending is responsible for the reduction in MeCP2 levels, we devised a simple strategy to edit the TGA stop codon found in CTD alleles to TGG (tryptophan) using an ABE (Figure 4A). In this case the next downstream stop codon would result in a –QGASX ending which is expected to be benign (Figure 4B). An advantage of our strategy is that the obligate presence of the pathogenic –PPX ending in all RTT CTD alleles means that a single sgRNA could be used to treat all mutations in the class, amounting to about 10% of all RTT cases.

**Figure 4.**
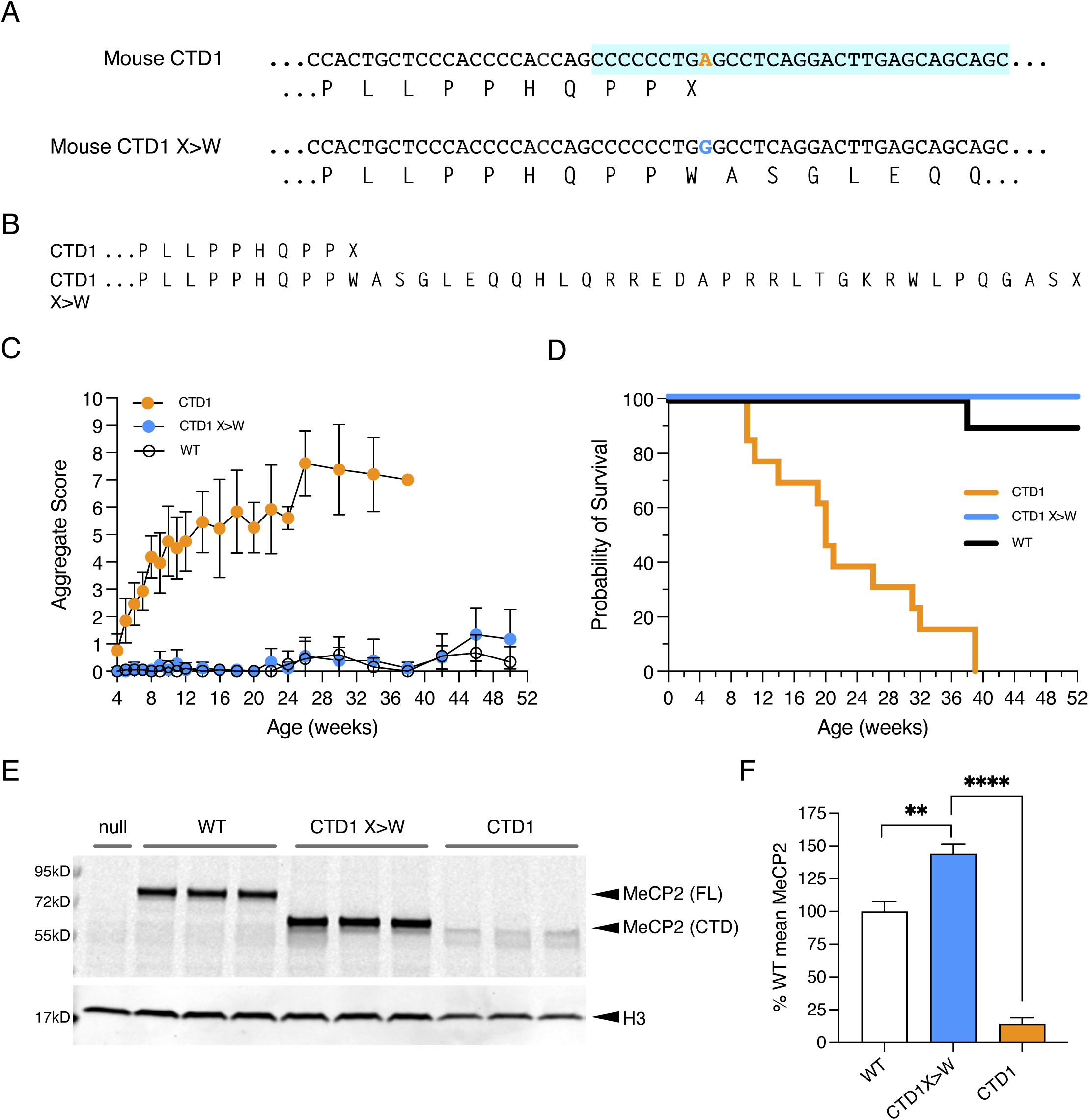
A CTD1 X>W knock-in mouse models adenine base editing of the CTD1 stop codon. (**A**) The genomic sequence of mouse CTD1 and CTD1 X>W alleles. The mutated adenine is shown in orange, and the sequence common to all mouse CTD alleles is shaded in turquoise. (**B**) The amino acid consequences of alleles in (A). (**C**) Phenotypic scoring of hemizygous male mice with CTD1 (n=13) and CTD1 X>W (n=9) knock-in alleles, and WT male littermates of CTD1 X>W animals (n=10). Mean +/− standard deviation (sd). (**D**) Kaplan-Meier plot of survival of animals shown in (C). (**E**) Western blot of whole brain protein from 6 week old male mice hemizygous for *Mecp2*-null, CTD1, CTD1 X>W and WT alleles. Full-length (FL) and C-terminally deleted (CTD) MeCP2 proteins are indicated. Histone H3 is used as a loading control. (**F**) Quantification of (E). N=3 per genotype, mean +/− sd. Unpaired two-tailed t-test: CTD1 X>W vs CTD1 P<0.0001 (****), CTD 1 X>W vs WT P=0.0021 (**).

Conversion of A to G in the stop codon will install a tryptophan codon in its place. To test the consequences of this replacement, we made a new CTD1 X>W knock-in allele in mouse embryonic stem cells (mESCs) that contained tryptophan at this position, thereby mimicking the outcome of the proposed edit (Fig. 4 SF1A, B). The truncated protein is slightly larger than MeCP2 CTD1 and CTD2 due to the extended missense tail (Figure 4B) and we found that ESC-derived neurons carrying this CTD1 X>W allele restored the CTD MeCP2 protein to WT levels (Fig. 4 SF2A). To demonstrate that this approach could be applied to multiple CTD alleles, we generated a corresponding CTD2hu X>W allele and again observed recovery of wildtype levels of MeCP2 protein in mESC-derived neurons (Fig. 4 SF2B). Importantly, CTD1 X>W hemizygous mice were indistinguishable from WT littermates, did not develop RTT-like phenotypes (Figure 4C, Fig. 4 SF3A) and survived to at least one year, in contrast to the original CTD1 mouse line (Figure 4D). As with the mESC-derived neurons, the X>W change restored MeCP2 protein and mRNA levels (Figure 4E, F, Fig. 4 SF3B). We conclude that introduction of a novel tryptophan and extended missense tail at this position has no discernible effect on protein function. In order to test the efficacy of base editing reagents to install the X>W allele, we next sought an experimental system that reproduced the relative MeCP2 protein levels produced from different mouse knock-in alleles. To express the protein at consistent physiological levels, we used the HEK cell-based Flp-In T-REx system (Invitrogen), previously found to perform better than transient transfections for NMD studies (Gerbracht et al., 2017). As MeCP2 is usually found at low levels in dividing cells and is only highly expressed in post-mitotic differentiated cells, tetracycline-inducible cDNA constructs were used so that transgene expression could be kept off during standard cell culture and induced for a limited period to examine mRNA and protein expression. Constructs based on the major e1 *MECP2* isoform were recombined into a single genomic locus using Flp recombinase to create an isogenic series of single-copy *MECP2* transgenic cell lines. CTD mutations replicating mouse alleles (CTD1, CTD2hu and CTD2mo) and patient mutations (c.1157_1197del, CTD1 and c.1164_1207del, CTD2) were introduced into mouse and human full-length cDNAs (Figure 5A, Fig. 5 SF1A) and integrated into the cells. Upon induction of transgene expression, these cell lines recapitulated the reduced MeCP2 protein and mRNA levels previously seen for the knock-in alleles (Figure 5B-D, Fig. 5 SF1B, C). The finding that the effect of the CTD mutations on protein and mRNA levels is well reproduced using expression constructs with a heterologous promoter, 3’ UTR and polyadenylation signal, and no intronic sequences, reinforces the notion that this is a post-transcriptional defect.

**Figure 5.**
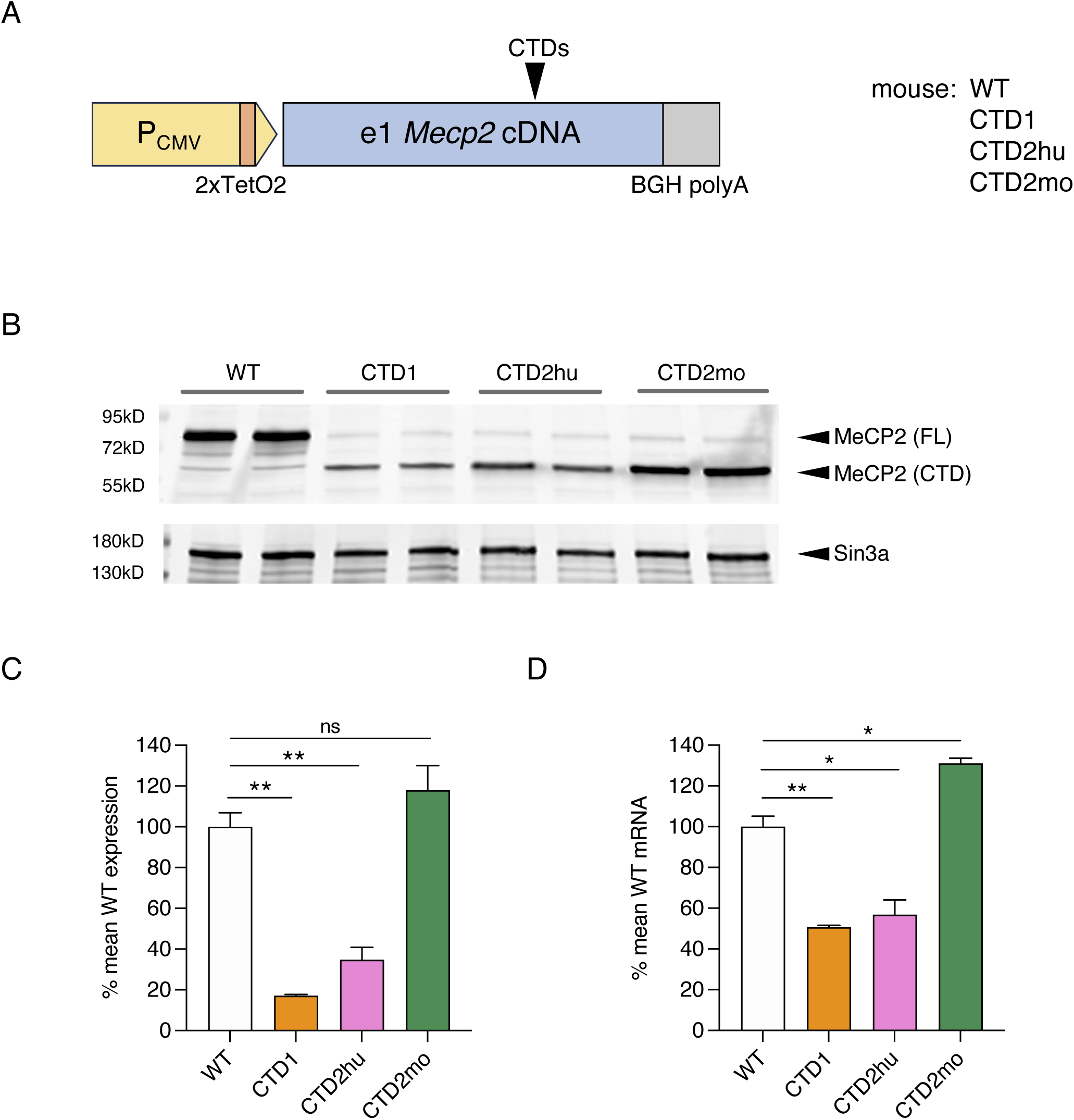
Flp-In T-REx cell lines reproduce the reduction in MeCP2 protein and mRNA seen with CTD knock-in mouse alleles. (**A**) Schematic of *Mecp2* transgenes in Flp-In T-REx cell lines. Deletions are introduced into a full-length e1 *Mecp2* cDNA, with a bovine growth hormone (BGH) polyadenylation signal and tetracycline-inducible CMV promoter. (**B**) Western blot with whole cell lysates from independent Flp-In T-REx clones carrying mouse cDNA transgenes (24 hours tetracycline induction). Sin3a loading control. (**C**) Quantification of MeCP2 protein expression from (B). N=2 clones per genotype, mean +/− sd. Unpaired two-tailed t-test: WT vs CTD1 P=0.0035 (**), WT vs CTD2hu P=0.0099 (**), WT vs CTD2mo P=0.2101 (ns). (**D**) Quantification of *Mecp2* transgene mRNA from the same experiment as (B) and (C). N=2 clones per genotype, mean +/− sd. Unpaired two-tailed t-test: WT vs CTD1 P=0.0057 (**), WT vs CTD2hu P=0.0208 (*), WT vs CTD2mo P=0.0171 (*).

The most commonly used ABEs consist of an evolved Tad deaminase domain(s) fused to a nicking *S. pyogenes* (Sp) Cas9, requiring an NGG PAM following the 20 nucleotide protospacer sequence (Gaudelli et al., 2017; Richter et al., 2020). Due to the cytosine-rich nature of the C-terminal deletion-prone region, there were no suitable PAMs available that would appropriately position the ABE to edit the target A (Figure 6A). Engineered SpCas9 PAM variant enzymes have been developed with relaxed PAM requirements to yield SpG that requires a single G PAM (NGN), and SpRY which has a relaxed sequence requirement but works most efficiently with either a single G or A (NG/AN) (Walton et al., 2020). We designed two gRNAs which would work with either SpG or SpRY ABEs (guides 1, A7 and 2, A4), and one with SpRY only (guide 5, A5) (Figure 6A). Plasmids encoding U6 promoter-expressed guide 1 or 2 and CMV promoter ABEmax-SpG (Walton et al., 2020) were transfected into T-REx cells with a mouse CTD1 (moCTD1) transgene, but no editing of an amplicon spanning the target site was detected by Sanger sequencing. As the two gRNAs position the target A7 at the edge of the optimal editing window for ABEmax (Gaudelli et al., 2017), next we tested ABE8e-SpG and ABE8e-SpRY constructs containing a further evolved TadA domain with increased editing efficiency and a wider editing window (Richter et al., 2020) (Figure 6B). These ABEs were able to edit the moCTD1 transgene with high efficiency using all gRNAs tested (Figure 6C), reaching 50-60% editing at the target A without selection of transfected cells. Editing with ABE8e-SpG was somewhat more efficient than with ABE8e-SpRY. Although editing at the target site was similarly efficient for all gRNAs, bystander editing at two nearby As (+6 and +9 relative to the target A) differed, with guide 1 giving minimal bystander editing, whereas using guides 2 and 5 resulted in significant bystander editing, particularly at the +6 position. Guide 1 extends furthest upstream (target A7), placing the bystander As furthest away from the editing window. Insertion or deletion mutations (indels) were low for all combinations (Fig. 6 SF1A).

**Figure 6.**
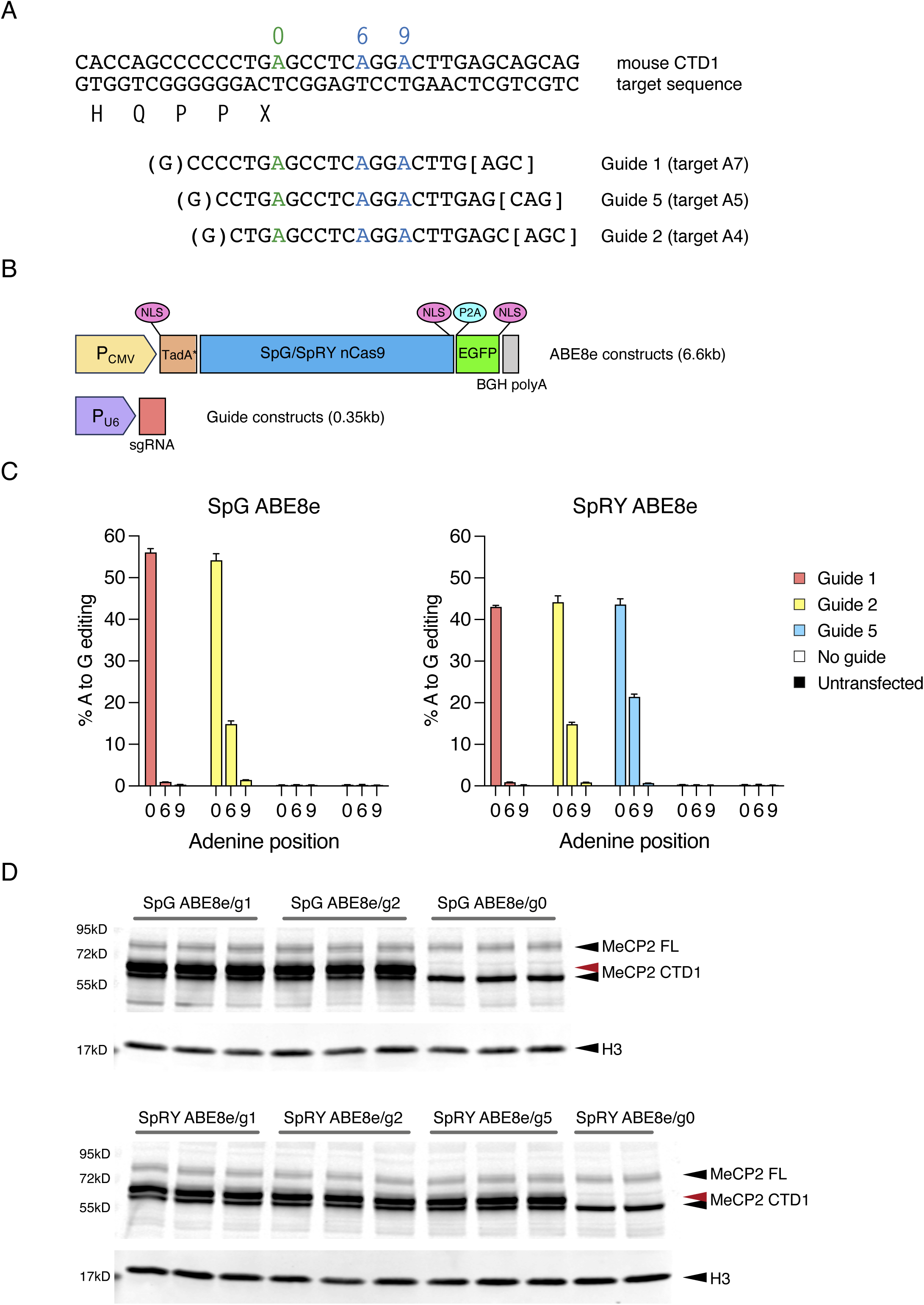
Base editing of mouse CTD transgenes in Flp-In T-REx cell lines. (**A**) Target genomic sequence and sgRNAs. The target A (position 0) is shown in green, with two bystander As within the guide sequence shown in blue (positions +6 and +9). Target site protospacer sequences are shown along with the PAM; the gRNA spacers are flanked by an additional G added to promote RNA polIII transcription. (**B**) ABE and gRNA constructs used for transfection experiments. (**C**) Editing efficiency following transfection of mouse CTD1 Flp-In T-REx cells with ABE8e-SpG or ABE8e-SpRY base editors and gRNA expression plasmids. Editing efficiency at the target and bystander As is quantified by amplicon sequencing (n=3 transfections per ABE/guide combination). (**D**) western blot showing MeCP2 protein levels from the experiment in (C) after 24 hours induction of transgene expression. The red arrow indicates MeCP2 CTD1 protein after editing (CTD1 X>W).

To determine whether editing of the moCTD1 allele led to an increase in MeCP2 protein, expression of the transgene was induced for 24 hours five days after transfection of the ABE constructs and guides. The significant editing seen in Figure 6C was accompanied by a concomitant increase in the amount of MeCP2, and a size shift due to the increased length of the missense tail (Figure 6D, Fig. 6 SF1B, C). We anticipated that a single gRNA could be applied to the editing of different CTD mutations. To test this, we used ABE8e-SpG guide 1 to edit a mouse CTD2hu allele in T-REx cells. Editing efficiency was indeed comparable to that of the CTD1 allele (Fig. 6 SF1D, E).

As a final proof-of-concept experiment, we asked whether these reagents also edited the human mutation. There is one nucleotide difference between mouse and human sequence in the guide region, so we tested human-specific gRNAs, guides 1 and 2, on a huCTD1 T-REx line (Fig. 6 SF2A). The human sequence-targeted gRNAs performed in a very similar way to their mouse equivalents (Fig. 6 SF2B, C), with the highest editing efficiency of almost 80% achieved using ABE8e-SpG and human guide 1. The mouse guide 1, with one mismatch to the human target, was also tested on the huCTD1 transgene, resulting in only a modest decrease in editing efficiency. We conclude that our DNA base editing strategy offers promise for future development as a therapeutic for RTT caused by CTD mutations.

## Discussion

The present study provides a simple genetic explanation for the initially puzzling observation that C-terminal deletions in the human *MECP2* gene cause RTT in some individuals but appear to be benign in others. In-frame deletions remove only a small number of amino acids from a region of MeCP2 that appears, from human genomic sequencing data, to be highly tolerant of mutation. CTDs causing shifts to the +1 and +2 frames both remove the entire C-terminus after the deletion site. Based on previous work in mice indicating that the detailed amino acid sequence close to newly created C-termini influences the level of MeCP2 expression, we compared the C-terminal sequences in databases of RTT patients and the general population. It emerged that RTT truncations terminated with the codons –PPX, whereas apparently benign truncations terminated in –SPRTX (Figure 7A). Using mouse models, we verified that the two prolines prior to the stop codon are essential contributors to the resulting severe deficiency of MeCP2 protein (Figure 7B). Cell lines containing CTD transgenes replicated these findings and were used to show that DNA base editing of the stop codon following the two prolines leads to readthrough to the next stop codon, preventing loss of MeCP2 protein (Figure 7C). Such genome editing could potentially be used to treat RTT patients with CTD mutations.

**Figure 7.**
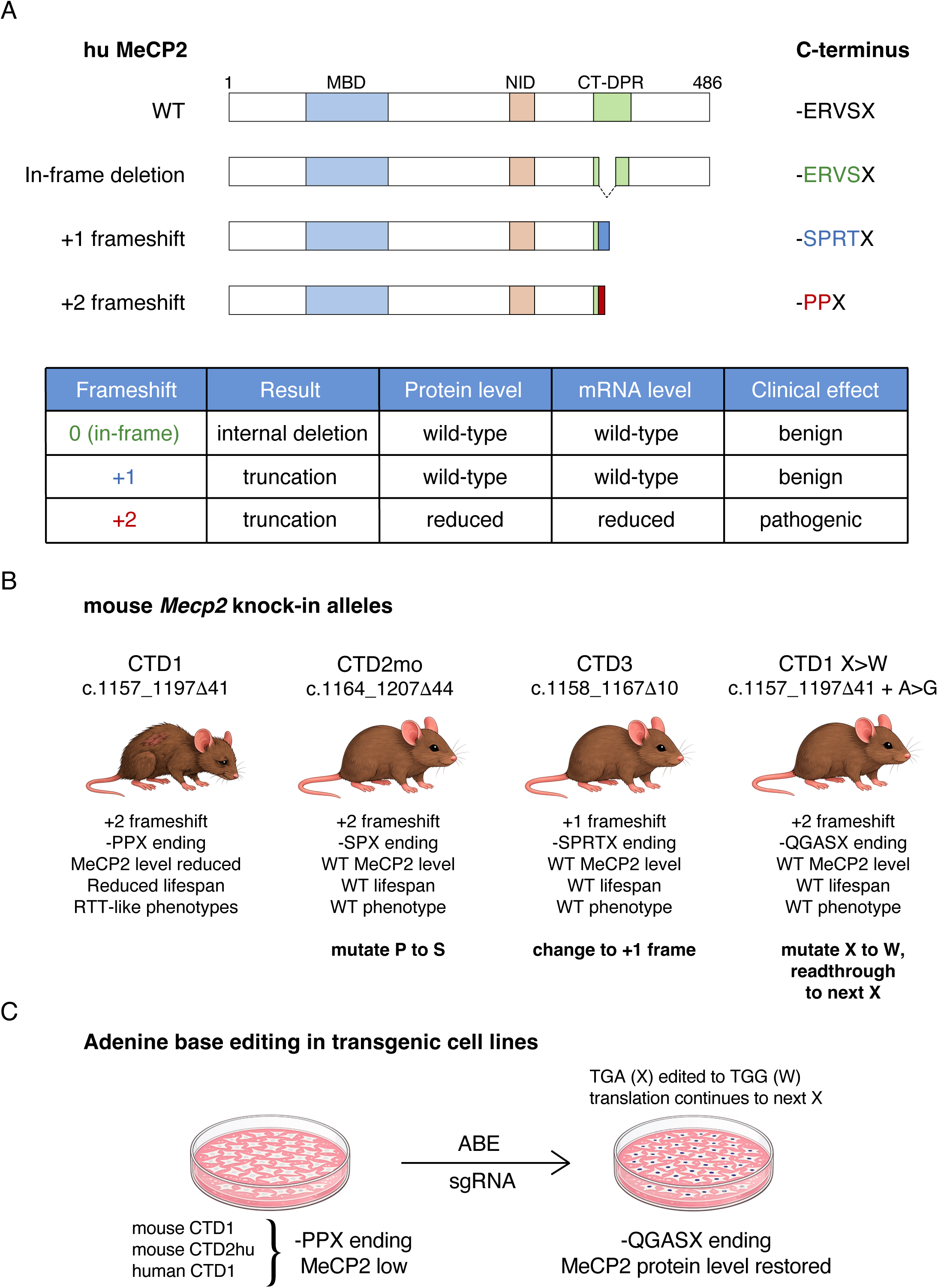
Summary of CTD alleles used in this study. (**A**) Human WT MeCP2 protein and the effect of the three possible reading frames remaining after deletions in the CT-DPR (**B**) *Mecp2* knock-in mouse models. The phenotype of the hemizygous CTD1 mouse can be ameliorated by altering the C-terminal amino acid sequence in several ways. All of these lead to non-PPX endings. (**C**) Efficient X>W editing of CTD transgenes using ABEs and sgRNAs in transfected cells. This replicates the CTD1 X>W allele and demonstrates the potential for therapeutic use of adenine base editing for RTT patients with CTDs.

The reason why mutations in the disordered C-terminal domain of MeCP2 should cause RTT has been the subject of speculation (Cheng et al., 2014; Li et al., 2020). This part of the protein is remote from the well characterised MBD and NID domains of MeCP2, and mouse models in which this entire region of the protein is missing have no RTT-like phenotype (Tillotson et al., 2017), suggesting that it is dispensable. Our previous work with mouse models of CTD mutations made clear that the truncations themselves do not interfere significantly with MeCP2 function. In particular, CTD2 mice in which the –PPX terminus is replaced by –SPX are like wildtype littermates despite persistence of the severe truncation. We show here that it is greatly reduced abundance of truncated MeCP2, presumably triggered by translational stalling at the –PPX codons, that drives the RTT phenotype.

A likely explanation for MeCP2 deficiency in CTD mutants is that the combination of consecutive prolines juxtaposed with a stop codon interferes with translation by the ribosome. Runs of multiple prolines are known to be problematic for translational elongation, likely due to the atypical geometry of this amino acid, with the translation factors eIF5A in mammals and EF-P in bacteria playing an important role in facilitating their translation (Gutierrez et al., 2013; Huter et al., 2017; Schuller et al., 2017). The effect of multiple prolines on translational termination is less well understood, although there is evidence from bacteria and eukaryotic cells that they may hinder efficient termination (Hayes et al., 2002; Janzen et al., 2002; Matheisl et al., 2015). In *E. coli* certain C-terminal sequences can trigger SsrA tagging, whereby the nascent polypeptide and translating RNA are targeted for degradation, thereby rescuing the stalled ribosome (Hayes et al., 2002). Interestingly, a sequential pair of prolines prior to the stop codon showed the strongest effect and, whereas seven *E. coli* proteins ending in PPX would be expected by chance, there are in fact none. In human cells, translation of a stalling peptide from human cytomegalovirus (hCMV) was used to trap the translational termination complex for structural studies. This effect was found to be dependent on the presence of two proline residues at the C-terminus of the peptide and the translational termination factor eRF1 (Matheisl et al., 2015).

Our findings have important clinical implications. We have shown here a contemporary pedigree containing several individuals with the frameshifting C-terminal deletion c.1159_1210, p.P387SX5. The CTD mutation was first identified after prenatal screening of a female foetus and was assigned as disease-causing, with the family informed that she was likely to develop Rett syndrome. After further investigation the same mutation was found in several healthy family members, including the child’s father, and the family were reassured that the mutation was likely to be benign due to the +1 frameshift and –SPRTX ending. Our results promise to resolve the discrepancy between genetic inference and clinical manifestation in these cases. Genetic testing can now incorporate this information to provide reassurance in cases where the pathogenic –PPX terminus is absent. The outcome of deletions in the CT-DPR can be predicted based on the exact location and size of the deletion (Figure 8). On a cautionary note, despite clinical examples arguing that these are phenotypically neutral mutations, it will be wise to maintain vigilance. While benign CTDs will not give rise to RTT, the possibility that less severe consequences accompany these mutations deserves future scrutiny.

**Figure 8.**
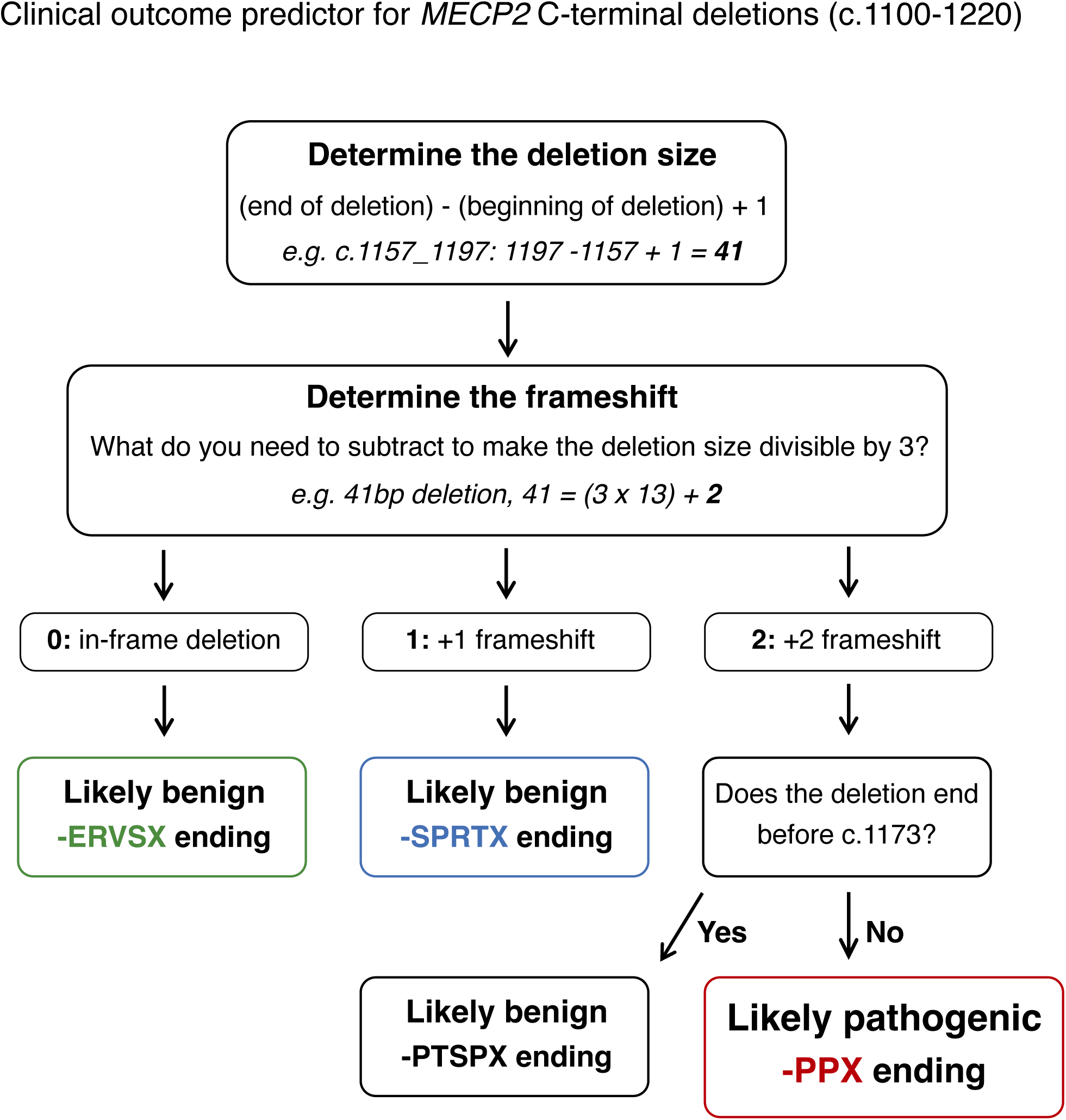
Flow chart for predicting the likely clinical prognosis of deletions in the CT-DPR of human *MECP2* based on the findings in this study.

The shared nucleotide sequence features of pathogenic CTDs raise the possibility that genome editing can be therapeutically beneficial. Using an SpCas9-based DNA base editor, we have shown that one guide RNA is potentially capable of targeting all pathogenic CTD mutations with the –PPX ending and can edit CTD transgenes efficiently in human cell lines. The edits replace the –PPX terminal sequence by –PPW– and add a short stretch of out-of-frame amino acid sequence before encountering the next stop codon. Modelling in mice indicates that the introduction of tryptophan plus downstream amino acids completely rescues the effects of the pathogenic mutation. Next steps will involve testing for phenotypic rescue after DNA base editing in an animal model, and addressing the issue of AAV-mediated delivery, which is relatively inefficient with most currently available vectors. The ultimate goal is to provide a unified therapeutic approach to this varied but unexpectedly homogeneous category of RTT-causing mutations.

## Materials and Methods

### Human mutation data

RTT patient mutation data was taken from RettBASE (Christodoulou et al., 2003), formerly hosted at http://mecp2.chw.edu.au/ and downloaded 17/08/2017 (Supplementary Table 1). The data held in RettBASE has now been incorporated into ClinVar (RRID:SCR_006169) in an abbreviated form (https://www.ncbi.nlm.nih.gov/clinvar/submitters/504290/). High confidence RettBASE CTD alleles were those where at least one individual carrying that mutation had been diagnosed with classical Rett syndrome and carried a *de novo MECP2* mutation (i.e. not present in either unaffected parent).

GnomAD data was taken from release v4.1.0,RRID:SCR_014964, downloaded from https://gnomad.broadinstitute.org/news/2024-04-gnomad-v4-1/ on 21/11/2024 (Supplementary Tables 5-8). *MECP2* mutations throughout are numbered according to transcript ENST00000303391.11 and amino acids are numbered according to the e2 MeCP2 isoform, as commonly used for RTT patient mutations. High confidence gnomAD CTD alleles were defined as those where at least one hemizygous (male) individual carried that mutation.

The family with the c1159-1210 deletion reached out to HZ for advice and this led to a detailed assessment of the medical history and the recommendation for expanded genetic testing in other family members.

### AlphaMissense data

Mean pathogenicity scores at each amino acid position of human MeCP2 (e2 isoform) were obtained from AlphaMissense (https://alphamissense.hegelab.org) (Cheng et al., 2023). The mean value for a sliding window of five amino acids was plotted at each position.

### Generation of *Mecp2* CTD mutations in mouse ES cells

*Mecp2* CTD knock-in alleles were generated in male mouse ES cell line JU09 (A gift from Joe Mee, University of Edinburgh), derived from 129/Ola E14Tg2a cells (RRID:CVCL_3505). Cells were transfected with a CRISPR/Cas9 plasmid based on pX330-U6-Chimeric_BB-CBh-hSpCas9, a gift from Feng Zhang (http://n2t.net/addgene:42230; RRID:Addgene_42230) (Cong et al., 2013) containing a sgRNA guide with protospacer sequence CCCCTGAGCCTCAGGACTTG within the deleted region for each allele, along with a 190nt single stranded oligodeoxynucleotide (ssODN) containing the desired DNA sequence changes, and a puromycin resistance plasmid for transient selection of transfected cells. For each knock-in allele 4 x 10^5^ ES cells were transfected with 500ng CRISPR/Cas9 plasmid, 200ng pPGK-Puro-pA plasmid and 10pmol ssODN using Lipofectamine2000 (ThermoFisher). After 24 hours 0.8μg/ml puromycin selection was added and maintained for 48 hours. Selected cells were plated at clonal density without selection and clones were picked after approximately 10 days.

### ssODNs

CTD3:

AGCAGCAGTGCCTCCTCCCCACCTAAGAAGGAGCACCATCATCACCACCATCACTCAGAGTCCCCAAAGGCGCCCGTGCCACTGCTCCCACCCCTCCACCTGAGCCCGCGGGCTCCGAGGACCCCACCAGCCCCCCTGAGCCCCAGGACTTGAGCAGCAGCATCTGCAAAGAAGAGAAGATGCCCCGAGG

CTD1 X>W:

AGCAGCAGTGCCTCCTCCCCACCTAAGAAGGAGCACCATCATCACCACCATCACTCAGAGTCCCCAAAGGCCCCCGTGCCACTGCTCCCACCCCATCAGCCCCCCTGGGCCTCAGGACTTGAGCAGCAGCATCTGCAAAGAAGAGAAGATGCCCCGAGGAGGCTCACTGGAAAGCGATGGCTGCCCCAAG

CTD2 X>W:

AGCAGCAGTGCCTCCTCCCCACCTAAGAAGGAGCACCATCATCACCACCATCACTCAGAGTCCCCAAAGGCCCCCGTGCCACTGCTCCCACCCCTGCCCCCCTGGGCCTCAGGACTTGAGCAGCAGCATCTGCAAAGAAGAGAAGATGCCCCGAGGAGGCTCACTGGAAAGCGATGGCTGCCCCAAGGAG

Clones were screened for hemizygous knock-in alleles by PCR on genomic DNA (forward primer: AGCATCAGAAGGTGTTCAGG, reverse primer: CCATAGGCTGAGTCTTAGCTGG), followed by cutting with SacII (CTD3, site gained) or SacI (CTD1 X>W, CTD2 X>W, site lost), and sequencing. Correctly edited clones were karyotyped and mycoplasma tested before further use.

### Differentiation of mouse ES cells into neurons

Verified ES cell knock-in clones were differentiated into neurons. At least two independently targeted clones were used for each genotype. Differentiation was carried out using a 4-/4+ retinoic acid (RA) procedure as previously described (Bain et al., 1995; Li et al., 1998). Neural precursor cells were seeded onto 6cm dishes at a density of 1.5×10^5^ cells/cm^2^ and harvested after 7 days by scraping into ice cold phosphate-buffered saline (PBS), pelleting the cells and snap freezing.

### Establishment of Mecp2 mutant mouse lines

Wildtype C57BL/6J mice (RRID:IMSR_JAX:000664) were purchased from Charles River Laboratories. CTD3 and CTD1 NS mouse lines were generated by injection of knock-in ES cells into mouse C57BL/6J blastocysts using standard methods. Chimeric males were crossed with C57BL/6J females and germline transmission from the ES cells was identified by coat colour. All agouti female pups were obligate heterozygotes for the *Mecp2* mutant allele due to its location on the X chromosome. This (N=1) and subsequent generations were bred by crossing Mecp2-mutant heterozygote females with C57BL6/J WT males. The genotypes of mutant lines were confirmed by sequencing. Routine genotyping was performed on ear biopsies using custom assays developed by Transnetyx, details available on request.

Mice were bred in a specific-pathogen-free facility in individually ventilated cages, with wood chippings, tissue bedding and environmental enrichment and were given *ad libitum* access to food and water. Rooms were maintained on a 12-hour light/dark cycle at 45-65% humidity at 20-24°C. Procedures were carried out by certified persons, licensed by the UK Home Office and according to the Animals (Scientific Procedures) Act 1986 under project licences (PPLs) 60/4547 and PP4326006. Ethical review of the PPLs was carried out by the University of Edinburgh Animal Welfare and Ethical Review Board (AWERB).

### Mouse phenotyping

Cohorts of hemizygous mutant male mice and WT littermates (N=3 backcross to C57BL/6J) were weighed and scored for a range of RTT-like phenotypes (mobility, gait, hindlimb clasping, tremor, breathing and general condition) to give an aggregate score between 0 and 12 as previously described (Cheval et al., 2012; Guy et al., 2007). Cohorts of at least 9 mutants and 9 wild-type littermates were scored for each knock-in line, with higher numbers being used where available to allow for any losses unrelated to the mutation, such as fighting. Sample sizes were selected based on our previous studies of *Mecp2* mutant mice and practical considerations of animal availability; no formal power calculation was performed. Animals were allocated to experimental groups without formal randomisation, as group assignment was determined by genotype; however, littermates were used where possible to minimise confounding effects. Males which would by necessity have to be singly housed during the study period were excluded. Scoring was carried out blind to genotype and to previous scores. Animals receiving a maximum score of 2 for tremor, breathing or general condition or which had lost 20% of their body weight had reached the severity limit of the experiment according to the Home Office license and were humanely culled. These animals were counted as having ‘died’ for the purposes of survival data. Animals of any genotype which were culled for reasons not linked to the mutation, such as fighting with cage mates, were removed from survival plots at that point (censored data). Scoring was carried out for one year, from 4 to 50 weeks of age, starting weekly (4-12 weeks), then every two weeks (12-26 weeks) and then every four weeks (26-50 weeks).

### Preparation of RNA and quantitative RT-PCR

Total cellular RNA was prepared from mESC-derived *in vitro* differentiated neurons, Flp-In T-REx cell lines and mouse brain using TriReagent (Sigma), according to the manufacturer’s protocol. cDNA was prepared from 1μg total RNA per reaction using a QuantiTect kit (Qiagen) and amplified in a qPCR reaction using SensiMix SYBR and Fluoroscein Master Mix (Bioline) in a LightCycler 480 (Roche). Primers for qPCR were as follows:

Mouse *Mecp2* mRNA: Exon3 F – ACCTTGCCTGAAGGTTGGAC, Exon4 R – GCAATCAATTCTACTTTAGAGCGAAAA

Mouse *Mecp2* primary transcript: Intron1 F – ACATGGCCGACAGAGTGC, Intron1 R – GCACCTGAGGAAGCAAACC

Flp-In T-REX *Mecp2* transgene (mouse and human): moMe2TREx F – GAGAGGAGCCTGTGGACAGC, BGHpA R – CACAGTCGAGGCTGATCAGC

Cyclophilin A mRNA (control): CypA F – TCGAGCTCTGAGCACTGGAG, CypA R – CATTATGGCGTGTAAAGTCACCA

Samples were run in triplicate and the amount of Mecp2 cDNA or primary transcript was calculated for each biological replicate after correction of CT values for PCR efficiency (using a dilution series to plot a standard curve) and normalisation to the control transcript Cyclophilin A. Each biological replicate was normalised to the mean of all WT samples for that transcript.

### Western blotting

*In vitro* differentiated ES cell-derived neurons, Flp-In T-REx cells and whole brain samples were prepared for western blotting as previously described (Ross et al., 2016). Extracts were run on TGX 4-20% gradient gels (BioRad) loading extract equivalent to approximately 5 x 10^5^ nuclei per well. For quantification, samples were run on duplicate gels and transferred to nitrocellulose membrane overnight in the cold at 25V. Western blots were processed as described previously (Guy et al., 2018; Ross et al., 2016) using the following antibodies: anti-MeCP2 (N-terminus): mouse monoclonal Men-8, RRID:AB_477235 (Sigma), anti-NeuN: rabbit polyclonal ABN78, RRID:AB_10807945 (Millipore), anti-histone H3: rabbit polyclonal ab1791, RRID:AB_302613 (Abcam), anti mSin3a: rabbit polyclonal ab3479, RRID:AB_303922 (Abcam). Western blots were developed with IR-dye secondary antibodies (IRDye 800CW donkey anti-mouse, RRID:AB_621847 and IRDye 680LT donkey anti-rabbit, RRID:AB_10956166, LI-COR Biosciences) and scanned using a LI-COR Odyssey machine. Images were quantified using Image Studio Lite software version 6.0.0.28 (LI-COR Biosciences). The ratio of NeuN:histone H3 signals for each lane was used to check for equal neuronal *in vitro* differentiation and clones that did not differentiate well were discarded.

MeCP2 levels were normalised to the histone H3 (neurons, whole brain) or mSin3a signal (Flp-In-TREx cells) for each lane to compare the amount of MeCP2/nucleus between samples.

### Production and characterisation of Flp-In T-REx cells

Flp-In™ T-REx™ 293 cells (RRID:CVCL_U427) were obtained from Invitrogen and maintained according to the manufacturer’s instructions. Cells were maintained under appropriate selection and routinely tested for mycoplasma. Tetracycline-inducible MeCP2 expression constructs were made by cloning human and mouse e1 cDNAs into plasmid pcDNA5/FRT/TO, Cat#V6520920 (Invitrogen). These were then inserted into the FRT site of Flp-In™ T-REx™ 293 cells according to the manufacturer’s instructions. Single-cell clones were verified by sequencing and single integration of the plasmid into the FRT site was confirmed by PCR or Southern blotting. Transgene expression was induced by addition of 0.5μg/ml tetracycline to the growth medium for 24 hours prior to cell harvesting and snap freezing of cell pellets.

### Transfection with ABEs/guides and quantification of DNA base editing

Plasmids ABE8e-SpG and ABE8e-SpRY were derived from pCMV-T7-ABEmax(7.10)-SpG-P2A-EGFP (Addgene plasmid # 140002; http://n2t.net/addgene:140002; RRID:Addgene_140002) and pCMV-T7-ABEmax(7.10)-SpRY-P2A-EGFP (Addgene plasmid #140003; http://n2t.net/addgene:140003; RRID:Addgene_140003) (Walton et al., 2020) by exchanging the ABEmax adenine deaminase domain for ABE8e (Alves et al., 2024). A separate plasmid, pGuide, a gift from Kiran Musunuru (Addgene plasmid # 64711; http://n2t.net/addgene:64711; RRID:Addgene_64711) was used to express sgRNAs using a human U6 promoter (Ding et al., 2013). Protospacer sequences were cloned into a BbsI site in pGuide using standard methods.

Flp-In T-REx cells were transfected with an equimolar mixture of ABE and guide plasmids. “No guide” controls received ABE + pGuide plasmid with no protospacer sequence (g0, empty guide). Cells were plated at 4 x 10^5^ cells per well in 6-well plates. The next day each well was transfected with 1000ng plasmid DNA and 6μl Lipofectamine 2000. Five days after transfection transgene expression was induced with 0.5μg/ml tetracycline. After 24 hours each well was harvested by trypsinisation, and the cells were split and snap frozen as two pellets for preparation of genomic DNA and protein for western blotting.

Genomic DNA was prepared from cell pellets using a Puregene Cell kit (Qiagen). Editing was quantified by MiSeq sequencing of a 190bp PCR amplicon spanning the target site: primers Me2 F6 – AGAAGGAGCACCATCATCACC, Me2 R2 – CCATAGGCTGAGTCTTAGCTGG. Amplicons were purified using DNA Clean and Concentrator-5 columns (Zymo Research) and sequencing libraries were made using NEBNext Ultra II DNA library prep kit for Illumina and NEBNext Multiplex Oligos for Illumina (New England Biolabs). Libraries were sequenced using a MiSeq Reagent Nano Kit v2 (300 cycles) (Illumina). Sequencing data was analysed using CRISPResso2, https://crispresso2.pinellolab.org/submission, RRID:SCR_021939 (Clement et al., 2019).

## Statistical analysis

All statistical analysis was performed using GraphPad Prism version 11.0.0, RRID: SCR_001066 (GraphPad Software). Datasets were compared using Student’s t-tests (unpaired, two-tailed, with or without Welch’s correction for unequal variance, as appropriate).

## Study approval

All animal experiments were performed under United Kingdom Home Office project licences (PPLs 60/4547 and PP4326006).

## Data availability

Data from this work are available from the authors.

## Acknowledgements

We would like to thank Emma Allen and the University of Edinburgh Central Transgenic Core (CTC) for production of mouse lines by blastocyst injection, Amanda Morris for MiSeq sequencing, Matthew Lyst, Jim Selfridge and Dirk-Jan Kleinjan for critical reading of the manuscript and John Christodoulou for helpful revision suggestions. This work was supported by Wellcome (Investigator Award 222507/Z/21/Z to AB), the Rett Syndrome Research Trust (Grant 13669477_13669480 to AB and JG), National Institutes of Health grant DP2CA281401 (B.P.K.), and the Simons Initiative for the Developing Brain (SIDB). A.B is a member of SIDB at the University of Edinburgh.

---

**Figure 2, Supplementary Figure 1.**
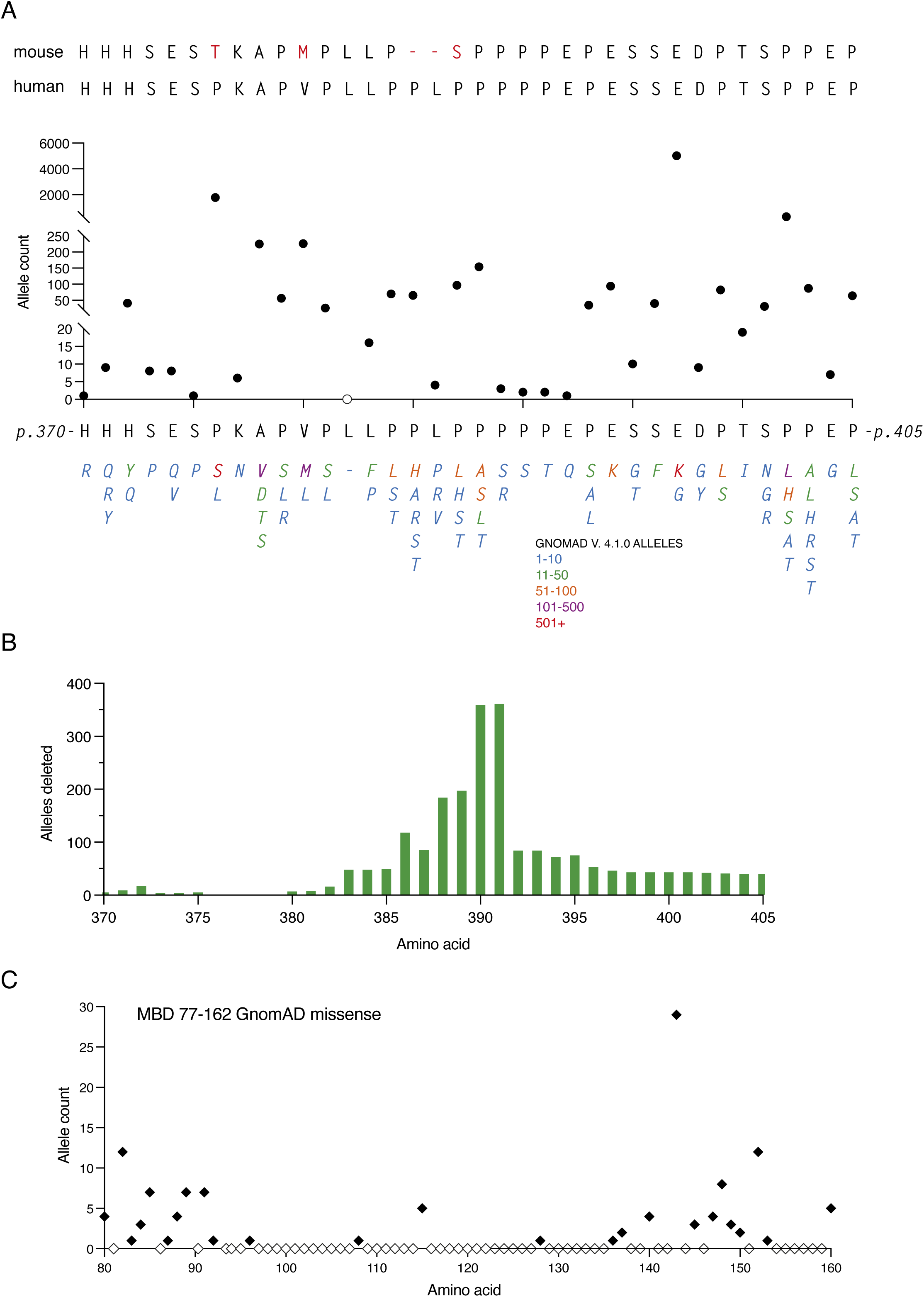
GnomAD v4.1.0 data from the CT-DPR. (**A**) GnomAD missense mutations. A comparison of mouse and human amino acid sequence in the CT-DPR, followed by a plot of the number of individuals in gnomAD with missense mutations present (filled circles) or absent (open circles) at each position. All amino acid changes found are listed below, colour-coded according to frequency. Hemizygous, heterozygous and homozygous mutations are included. (**B**) Plot of the total number of gnomAD alleles with in-frame deletions at each amino acid position in the CT-DPR (hemizygous and heterozygous individuals). (**C**) Plot of the gnomAD missense allele count in the MBD for comparison with (A). Mutations present (filled diamonds), no gnomAD changes (empty diamonds).

**Figure 2, Supplementary Figure 2.**
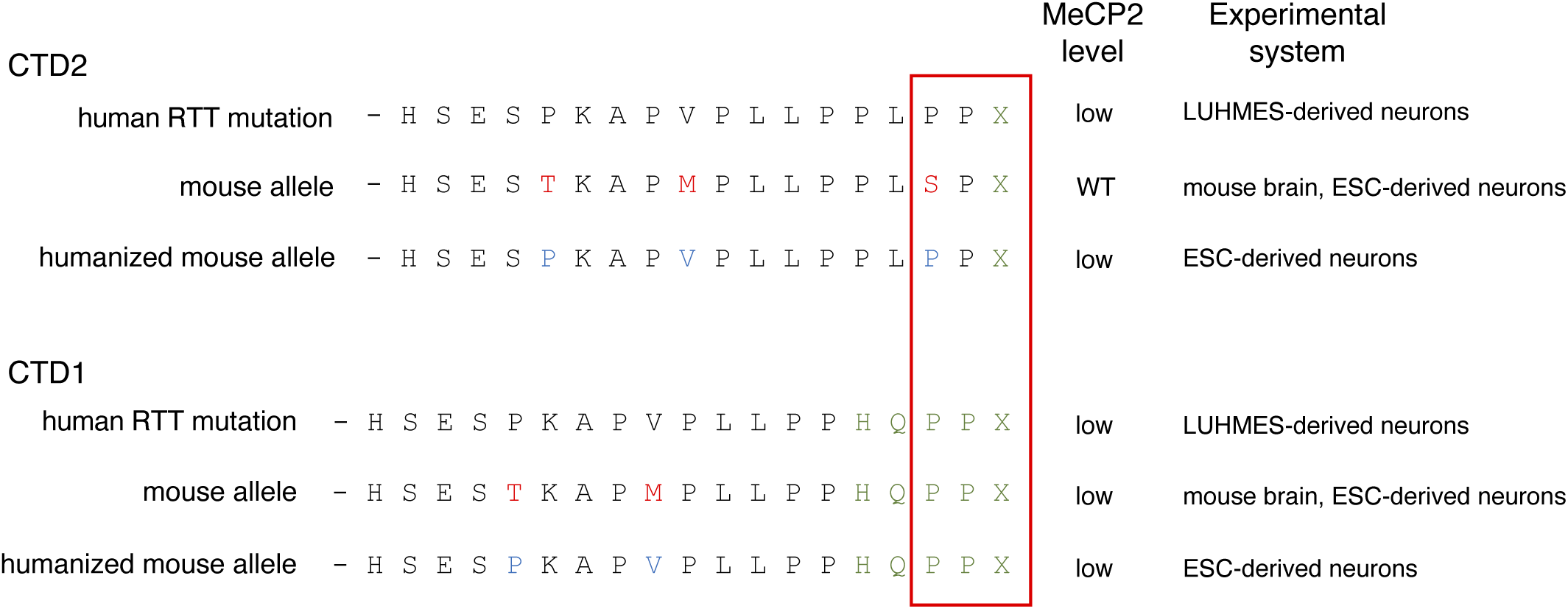
Summary of CTD C-terminal amino acid sequences and experimental MeCP2 protein levels for each.

**Figure 3, Supplementary Figure 1.**
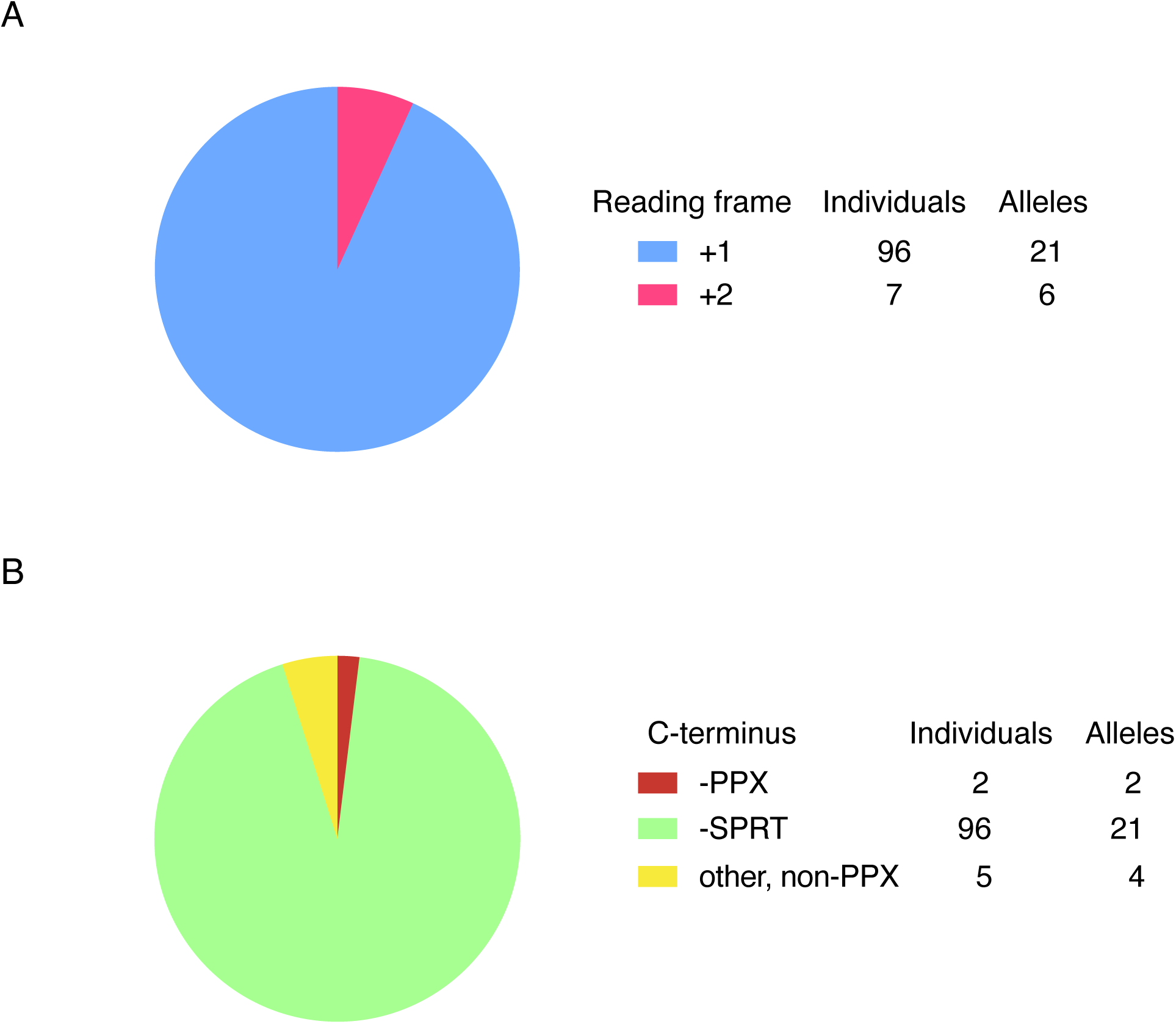
All frameshifting deletion alleles in gnomAD v4.1.0. (**A**) Number of individuals with each type of frameshift and number of different CTD alleles. (**B**) GnomAD entries shown in (A) classified by C-terminal amino acid sequence for individuals and CTD alleles.

**Figure 3, Supplementary Figure 2.**
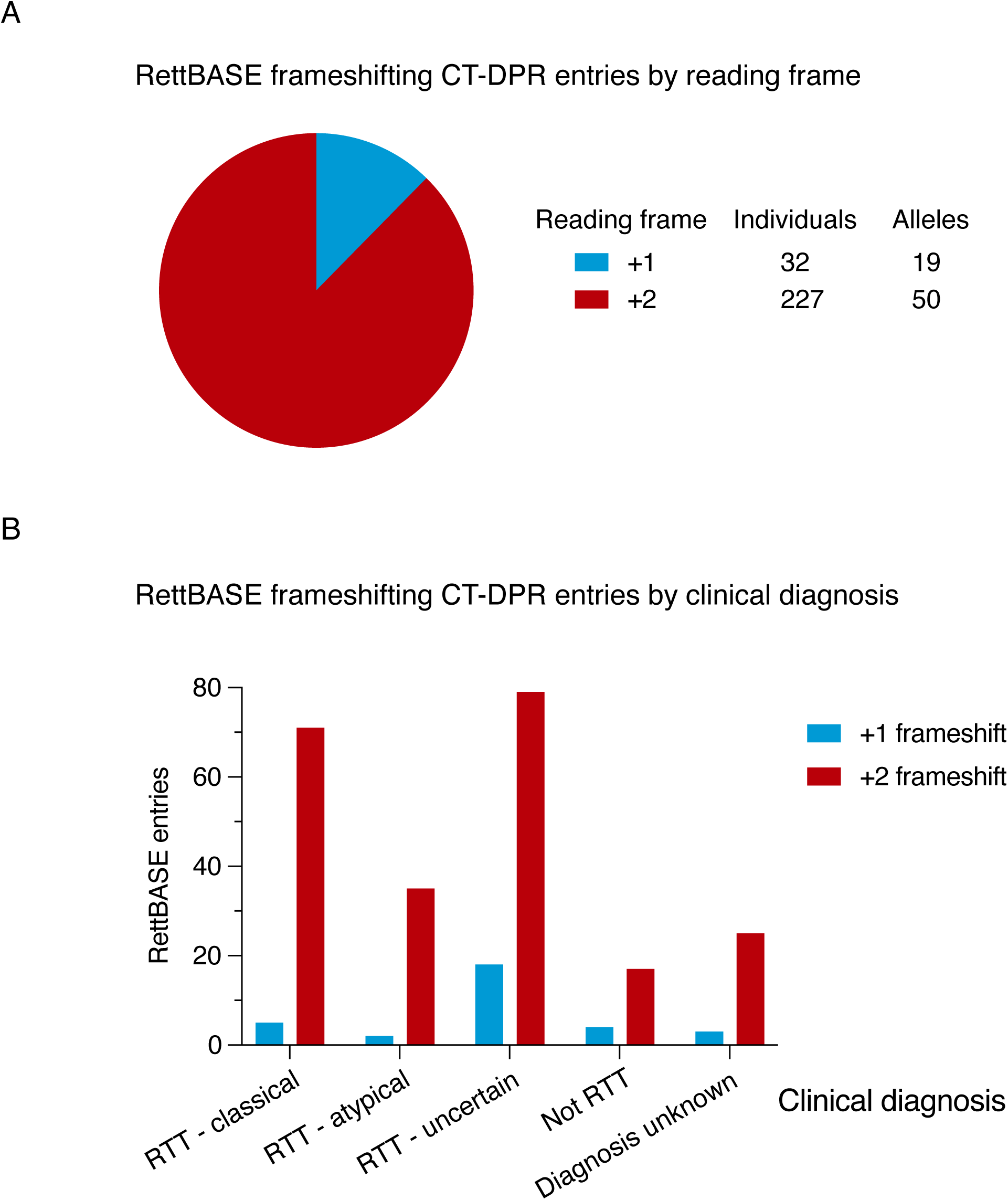
All frameshifting deletion alleles in RettBASE. (**A**) Number of individuals with each type of frameshift and number of different CTD alleles. (**B**) Entries shown in (A) grouped by clinical diagnosis displayed in RettBASE.

**Figure 3, Supplementary Figure 3.**
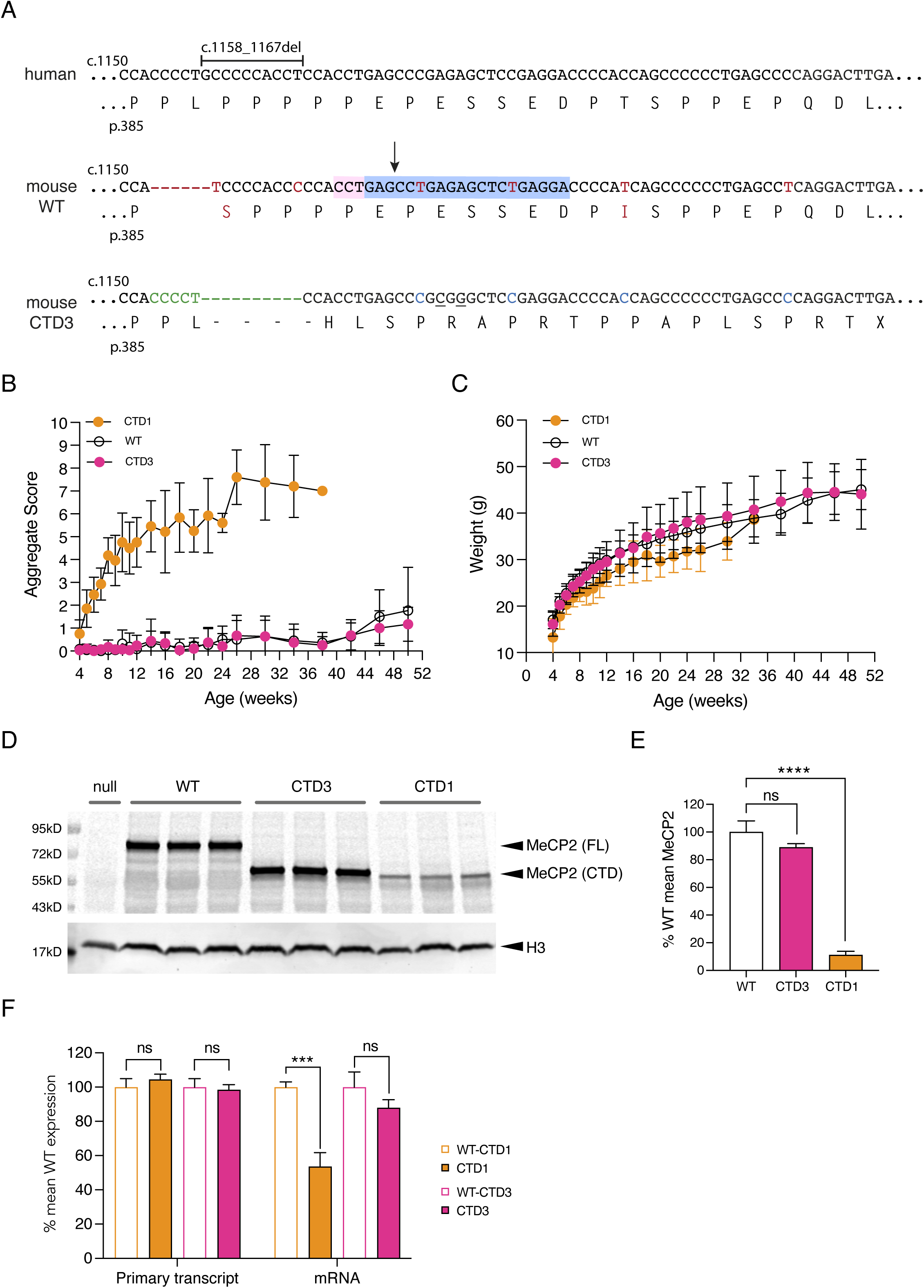
CTD3 mouse knock-in allele. (**A**) Human genomic and amino acid sequence showing location of CTD3 deletion (1158_1167del). Mouse WT genomic and amino acid sequences. Differences to human sequence are shown in red. The protospacer sequence (blue) and PAM sequence (pink) used to cut the WT allele for CRISPR editing are shown, with the cut site marked with an arrow. The mouse CTD3 knock-in allele is shown with nucleotide additions and deletions to the WT allele shown in green. Changes made to the mouse sequence to reproduce the human missense tail are in blue and two silent changes which introduce a diagnostic SacII site are underlined. (**B**) Phenotypic scoring of hemizygous male mice with CTD1 (n=13) and CTD3 (n=15) knock-in alleles, and WT male littermates of CTD3 animals (n=11). Mean +/− sd. (**C**) Body weights of animals shown in (B). Mean +/− sd. (**D**) Western blot of whole brain protein from 6 week old male mice hemizygous for *Mecp2*-null, CTD1, CTD3 and WT alleles. Full-length (FL) and C-terminally deleted (CTD) MeCP2 proteins are indicated. Histone H3 is used as a loading control. (**E**) Quantification of (D). N=3 per genotype, mean +/− sd. Unpaired two-tailed t-test: CTD3 vs WT P=0.079 (ns), CTD1 vs WT P<0.0001 (****). (**F**) Quantification of *Mecp2* primary transcript and mRNA in whole brain of 6 week old male mice as in (D). N=3 brains per genotype. Mean +/− sd. Unpaired two-tailed t-test: Primary transcript CTD3 vs WT littermates P=0.6687 (ns), CTD1 vs WT P=0.2585 (ns), mRNA CTD3 vs WT P=0.1055 (ns), CTD1 vs WT P=0.0008 (***).

**Figure 4, Supplementary Figure 1.**
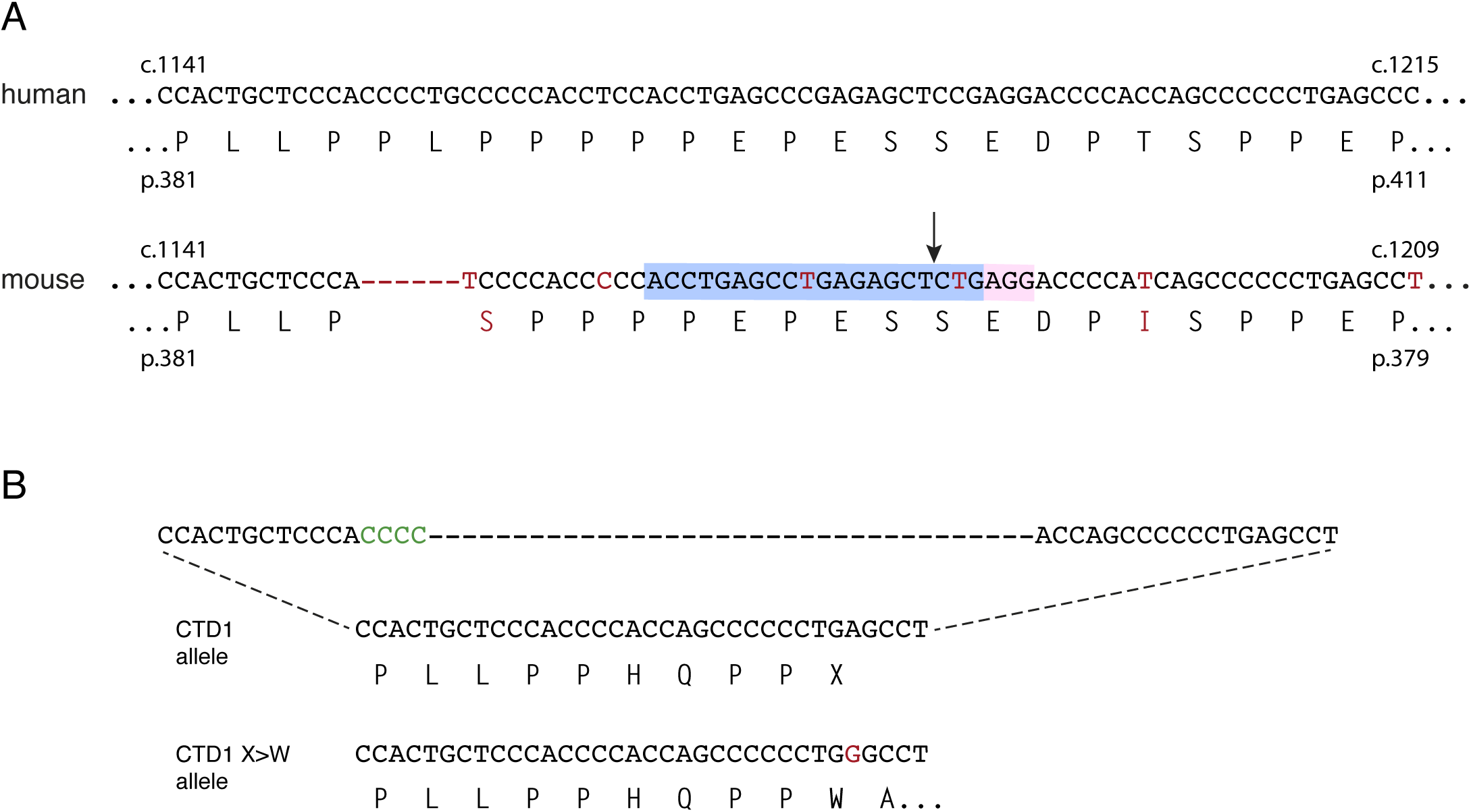
Structure of the mouse CTD1 X>W allele. (**A**) Human and mouse CT-DPR genomic and amino acid sequences. Differences between mouse and human sequence are shown in red. The protospacer sequence (blue) and PAM sequence (pink) used to cut the WT allele for CRISPR editing are shown, with the cut site marked with an arrow. (**B**) The mouse CTD1 knock-in allele compared to the CTD1 X>W allele. The single nucleotide A to G change is shown in red.

**Figure 4, Supplementary Figure 2.**
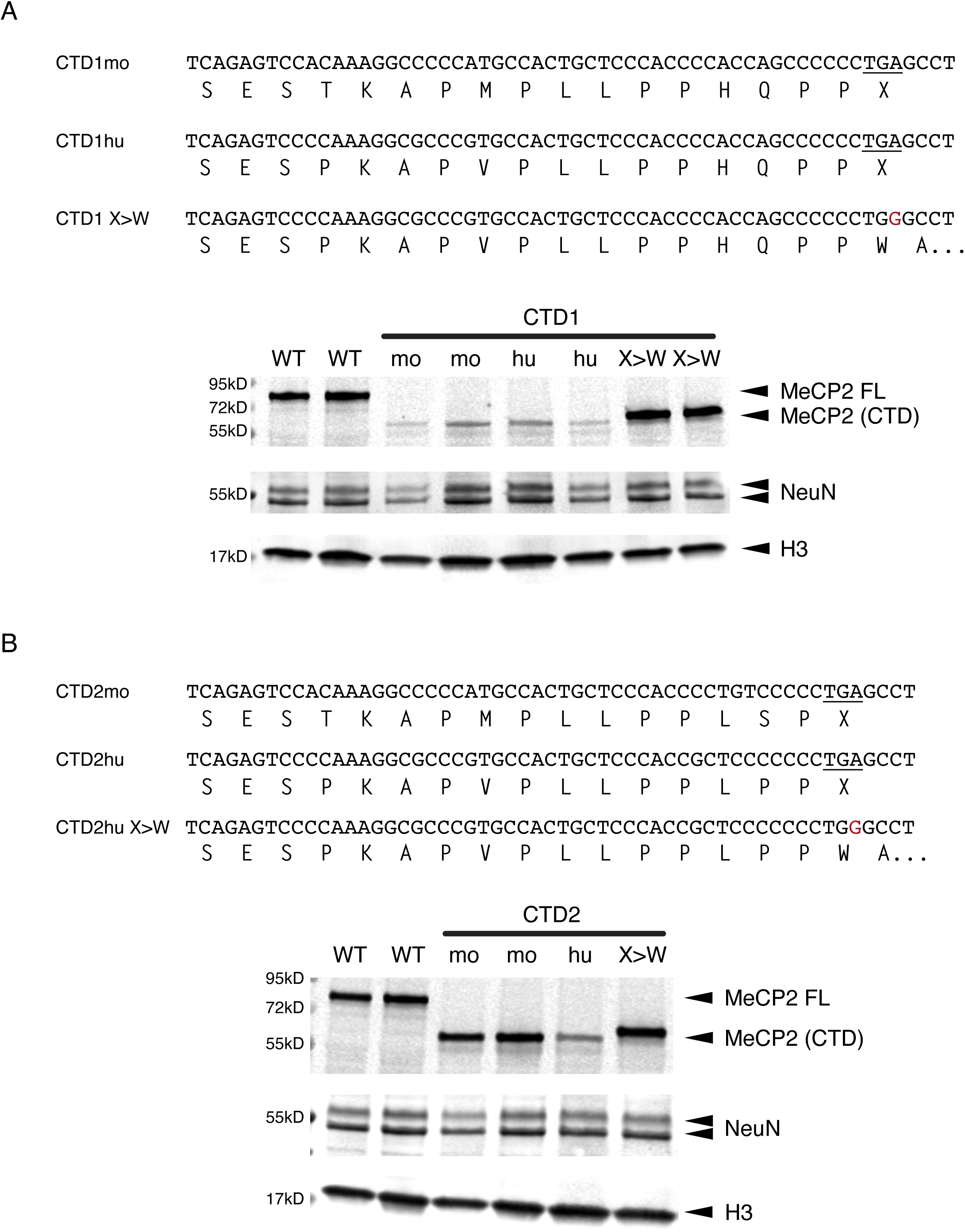
CTD mouse alleles: mESC-derived neurons. (**A**) genomic and amino acid sequences of 3 CTD1 mouse knock-in alleles and western blot of MeCP2 protein from mESC-derived neurons 7 days after plating neuronal progenitors. Two independent clones per genotype, histone H3 loading control, NeuN control for differentiation status. (**B**) genomic and amino acid sequences of 3 CTD2 mouse knock-in alleles and western blot of MeCP2 protein from mESC-derived neurons 7 days after plating neuronal progenitors.

**Figure 4, Supplementary Figure 3.**
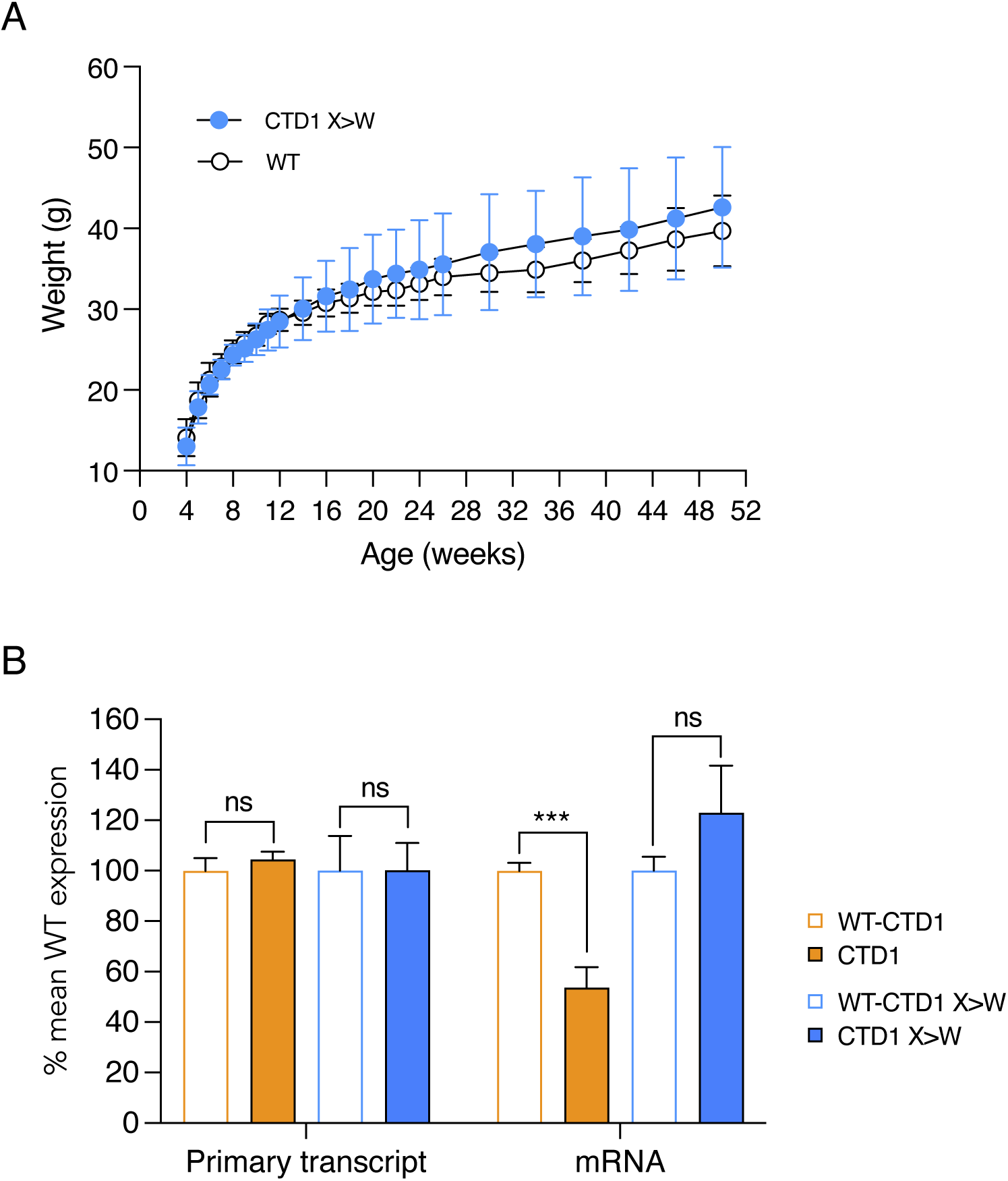
CTD1 X>W knock-in mice: weights and brain RNA levels. (**A**) Body weights of animals shown in Figure 4C and D. Mean +/− sd. CTD1 (n=13) and CTD1 X>W (n=9), WT male littermates of CTD1 X>W animals (n=10). (**B**) Quantification of *Mecp2* primary transcript and mRNA in whole brain of 6 week old male mice. N=3 brains per genotype. Mean +/− sd. Unpaired two-tailed t-test: Primary transcript CTD1 X>W vs WT littermates P>0.9999 (ns), CTD1 vs WT P=0.2585 (ns), mRNA CTD1 X>W vs WT P=0.1419 (ns), CTD1 vs WT P=0.0008 (***).

**Figure 5, Supplementary Figure 1.**
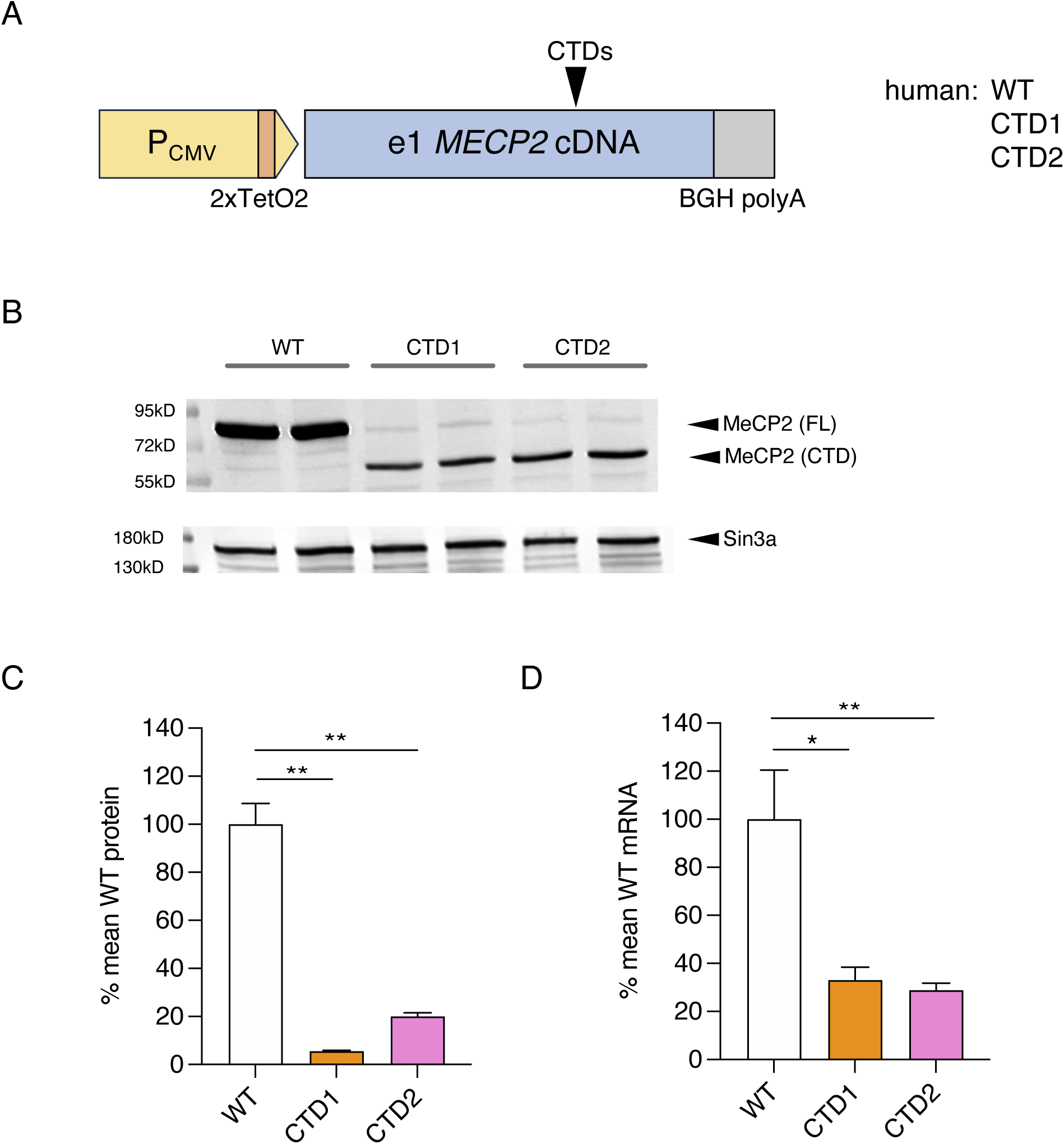
Flp-In T-REx cell lines with human *MECP2* transgenes. (**A**) Schematic of human *MECP2* transgenes in Flp-In T-REx cell lines. (**B**) Western blot with whole cell lysates from independent Flp-In T-REx clones carrying human cDNA transgenes (24 hours tetracycline induction). Sin3a loading control. (**C**) Quantification of MeCP2 protein expression from (B). N=2 clones per genotype, mean +/− sd. Unpaired two-tailed t-test: WT vs CTD1 P=0.0042 (**), WT vs CTD2 P=0.0060 (**). (**D**) Quantification of *Mecp2* transgene mRNA from the same experiment as (B) and (C). N=2 clones per genotype, mean +/− sd. Unpaired two-tailed t-test: WT vs CTD1 P=0.0126 (*), WT vs CTD2 P=0.0099 (**), WT vs CTD2mo P=0.0171 (*).

**Figure 6, Supplementary Figure 1.**
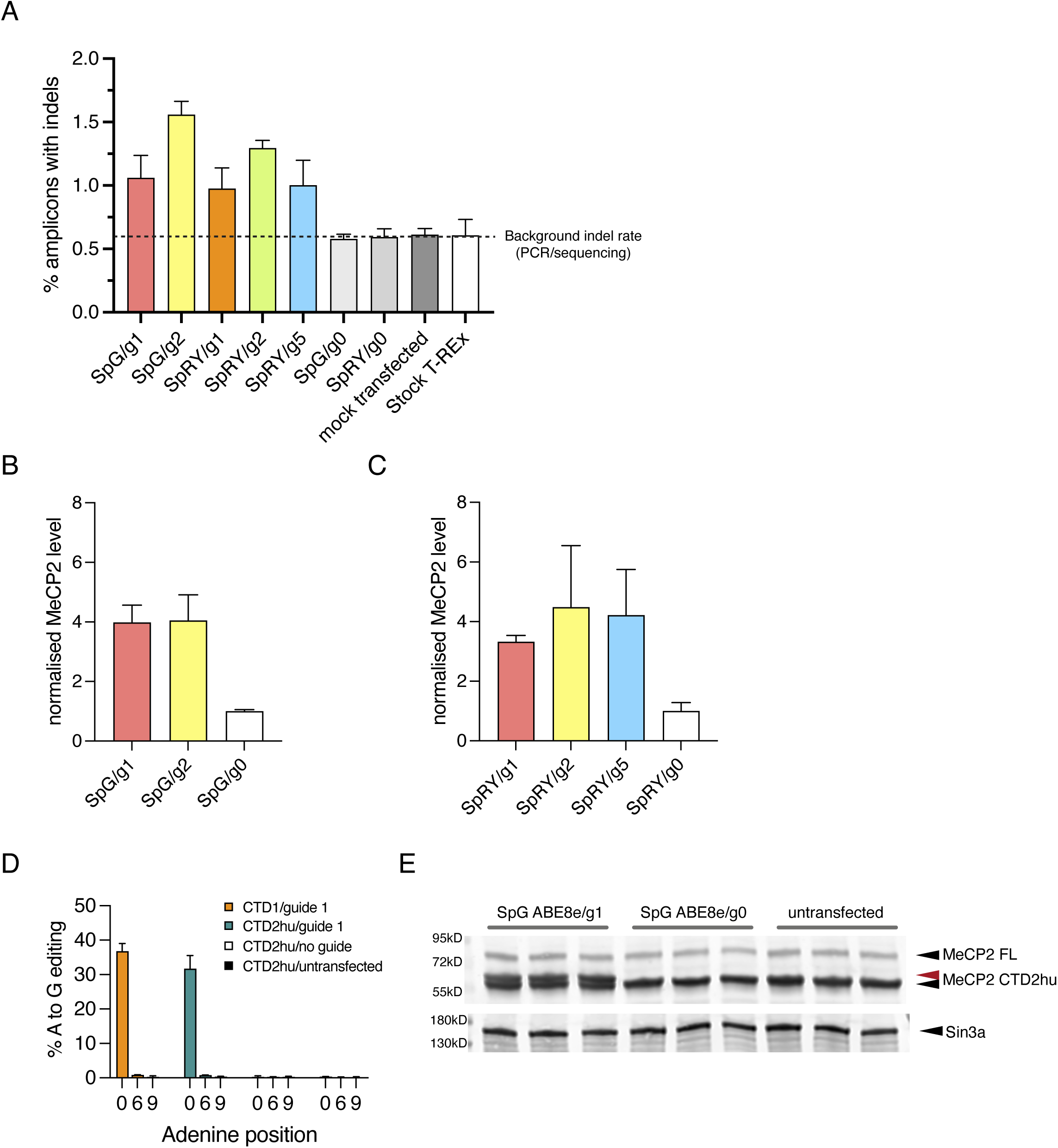
Base editing of CTD transgenes in Flp-In T-REx cell lines. (**A**) mouse CTD Flp-In T-REx cells treated with ABE/guide RNAs as shown in Figure 6C. Percentage of mapped reads with indels. Background indel rate is indicated based on the % indels found in amplicons from mock– or untransfected cells. (**B**) Quantification of total CTD MeCP2 levels from western blot in Figure 6D, SpG ABE8e. (**C**) Quantification of total CTD MeCP2 levels from western blot in Figure 6D, SpRY ABE8e. (**D**) Editing efficiency of SpG ABE8e/mouse guide 1 at target and bystander As in mouse CTD1 and CTD2hu Flp-In T-REx cells. Mean +/− sd, n=3 transfections per ABE/guide combination. (**E**) western blot showing MeCP2 protein levels from the mouse CTD2hu cells in (D) after 24 hours induction of transgene expression. The red arrow indicates MeCP2 CTD1 protein after editing (CTD1 X>W).

**Figure 6, Supplementary Figure 2.**
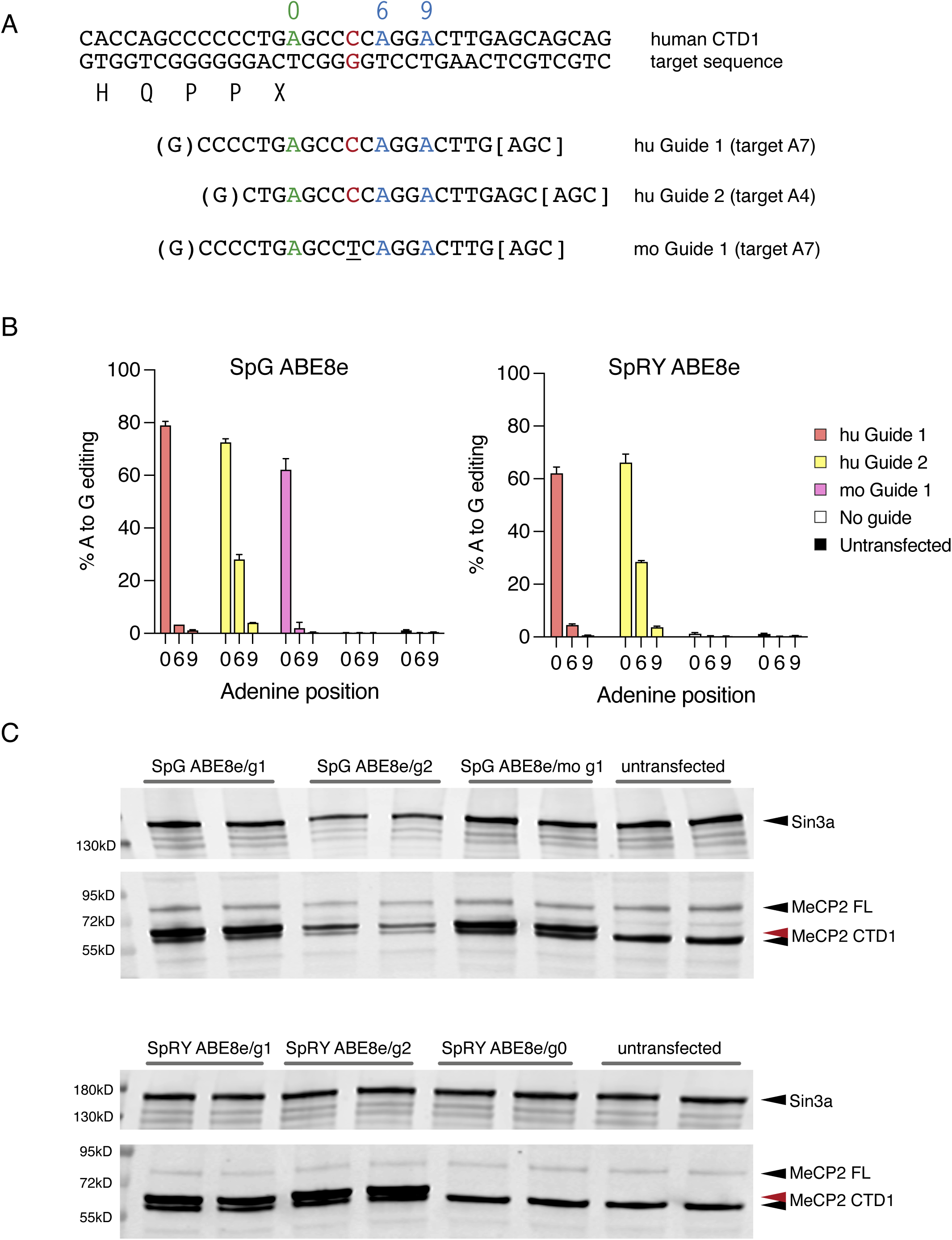
Editing of human *MECP2* transgenes in Flp-In T-REx cells. (**A**) Target genomic sequence and sgRNAs. Target A (position 0) is shown in green, with two bystander As within the guide sequence shown in blue (positions +6 and +9). The difference between human (red) and mouse (underlined) sequences is indicated. (**B**) Editing efficiency following transfection of human CTD1 Flp-In T-REx cells with ABE8e-SpG or SpRY base editors and guide RNA plasmids shown in (A). Editing efficiency at the target and bystander As is quantified by amplicon sequencing (n=3 transfections per ABE/guide combination). (**C**) western blot showing MeCP2 protein levels from the experiment in (B) after 24 hours induction of transgene expression. The red arrow indicates MeCP2 CTD1 protein after editing (CTD1 X>W). Sin3a loading control.

## Supplementary Table Legends

**Supplementary Table 1** All *MECP2* mutations held in RettBASE listed by location of first nucleotide change (4664 entries). Data downloaded 17/08/2017.

**Supplementary Table 2** RettBASE frameshifting mutations in the CT-DPR (c.1110-1210). All entries, listed by start of frameshift.

**Supplementary Table 3** RettBASE frameshifting alleles in the CT-DPR (c.1110-1210) listed by start of frameshift. The number of individuals with each allele is shown. Entries have been annotated with further details from RettBASE which were used to select a high confidence set of RTT CTD alleles (highlighted in pink): at least one individual with a diagnosis of Classical Rett syndrome and at least one individual with evidence for a *de novo MECP2* mutation. Frameshifts caused by insertions are also shown, with high confidence mutations highlighted in lilac. Alleles which are also found in gnomAD are indicated.

**Supplementary Table 4** Set of 15 high confidence RTT CTD alleles selected from RettBASE.

**Supplementary Table 5** gnomAD *MECP2* missense mutations in the CT-DPR listed by location. gnomAD release v4.1.0 downloaded 21/11/2024, nucleotide numbering according to transcript ENST00000303391.11.

**Supplementary Table 6** gnomAD *MECP2* in-frame deletions in the CT-DPR. High confidence mutations found in at least one hemizygous individual are highlighted in green.

**Supplementary Table 7** gnomAD *MECP2* missense mutations in the MBD listed by location (c.239-479).

**Supplementary Table 8** gnomAD *MECP2* frameshifting deletion alleles in the CT-DPR. Nine high confidence alleles which are present in at least one hemizygous individual are highlighted in blue. Two single mutations with a –PPX ending are highlighted in yellow.

